# SyFi: generating and using sequence fingerprints to distinguish SynCom isolates

**DOI:** 10.1101/2025.02.27.640502

**Authors:** Gijs Selten, Adrián Gómez-Repollés, Florian Lamouche, Simona Radutoiu, Ronnie de Jonge

## Abstract

The plant root microbiome is a complex community shaped by interactions among bacteria, the plant host, and the environment. Synthetic community (SynCom) experiments help disentangle these interactions by inoculating host plants with a representative set of culturable microbial isolates from the natural root microbiome. Studying these simplified communities provides valuable insights into microbiome assembly and function. However, as SynComs become increasingly complex to better represent natural communities, bioinformatics challenges arise. Specifically, accurately identifying, and quantifying SynCom members based on, for example, *16S rRNA* amplicon sequencing becomes more difficult due to the high similarity of the target amplicon, limiting downstream interpretations. Here, we present SynCom Fingerprinting (SyFi), a bioinformatics workflow designed to improve the resolution and accuracy of SynCom member identification. SyFi consists of three modules: the first module constructs a genomic fingerprint for each SynCom member based on its genome sequence, accounting for both copy number and sequence variation in the target gene. The second module then extracts a specific region from this genomic fingerprint to create a secondary fingerprint focused on the target amplicon. The third module uses these fingerprints as a reference to perform pseudoalignment-based quantification of SynCom member abundance from amplicon sequencing reads. We demonstrate that SyFi outperforms standard amplicon analysis by leveraging natural intragenomic variation, enabling more precise differentiation of closely related SynCom members. As a result, SyFi enhances the reliability of microbiome experiments using complex SynComs, which more accurately reflect natural communities. This improved resolution is essential for advancing our understanding of the root microbiome and its impact on plant health and productivity in agricultural and ecological settings. SyFi is available at https://github.com/adriangeerre/SyFi.

**Impact statement:** SyFi represents a significant advancement in microbiome research by enhancing the accuracy and resolution of synthetic community (SynCom) member identification. By leveraging natural intragenomic variation, SyFi improves the differentiation of closely related microbial strains, addressing a key challenge in amplicon-based sequencing analysis. This increased precision allows researchers to more reliably track microbial dynamics in complex SynCom experiments, leading to deeper insights into microbiome assembly, function, and host-microbe interactions. As a result, SyFi strengthens the interpretability of microbiome studies, ultimately contributing to a better understanding of plant health and productivity in both agricultural and ecological contexts.

**Data Summary:** The data reported in this article have been deposited in the National Center for Biotechnology Information Short Read Archive BioProject database. SyFi fingerprint generation was run on a collection of 737 human gut-derived bacterial genomes from Forster *et al*. (2019) (Genomic read data deposited in the ENA under project numbers ERP105624 and ERP012217) and 447 Arabidopsis-derived bacterial genomes (Selten *et al*., 2024b) (NCBI Project numbers PRJNA1138681, PRJNA1139421 (Genomes), and PRJNA1131834 (Genomic reads)). A list of the closed genomes used for SyFi validation can be found at https://github.com/adriangeerre/SyFi.

Subsequently, SyFi was validated on a complex SynCom dataset by pseudoaligning *16S rRNA* V3-V4 and V5-V7 amplicon reads (PRJNA1191388) to SyFi-generated fingerprints and comparing this to shotgun metagenomics-sequenced dataset of the same samples in Selten *et al*. (2024a) (PRJNA1131994). This complex SynCom dataset included the inoculation of the 447 bacterial isolates on Arabidopsis, Barley, and Lotus roots.

## Introduction

Sequencing the conserved *16S* ribosomal RNA gene (hereafter *16S rRNA*) is the benchmark method for microbiome community profiling across ecosystems (Jovel et al., 2016; Lucaciu et al., 2019; Parada et al., 2016; Weinroth et al., 2022). The *16S rRNA* gene is found in all prokaryotes and consists of intermixed variable and conserved regions. The conserved regions facilitate the design of universal primer sequences compatible with diverse prokaryotes, enabling the amplification of variable regions. These variable regions are taxonomically informative, allowing for the identification, classification, and quantification of prokaryotic micro-organisms within a community, thereby providing a broad overview of the microbiome composition across taxonomic levels (Clarridge, 2004). Compared to shotgun metagenome sequencing, amplicon sequencing is cheaper, faster, and applicable to host DNA- contaminated samples, though it has limitations (Liu et al., 2021). Most notably, it is prone to PCR-induced amplification biases and has limited taxonomic resolution, often restricted to the genus level (Liu et al., 2021).

Adding another layer of complexity, bacterial strains can harbor multiple copies of the *16S rRNA* gene, which may differ in sequence (Espejo & Plaza, 2018; Johnson et al., 2019; Lavrinienko et al., 2021; Louca et al., 2018; Schloss, 2021; Sun et al., 2013; Větrovský & Baldrian, 2013). Despite the fact that *16S rRNA* copy numbers can range from 1 to 15 copies per genome (Klappenbach, 2001), they are often not accounted for in microbiome analyses (Kembel et al., 2012). Moreover, intragenomic *16S rRNA* copies can differ due to biological variations such as insertions, deletions, or single nucleotide polymorphisms (SNPs), making them difficult to distinguish from sequencing errors (Pei et al., 2010; Schloss, 2021). This inherent biological variation can distort bacterial abundance estimates, leading to a skewed representation of microbiome composition (Schloss, 2021).

In recent years, synthetic community (SynCom) reconstitution experiments have emerged as a powerful approach in microbiome research (Jennings & Clavel, 2024; van Leeuwen et al., 2023; Vorholt et al., 2017). In this bottom-up approach, hosts are inoculated with simplified, yet representative microbial communities reconstituted from cultured bacterial isolates. SynCom experiments facilitate the investigation of host-microbe and microbe-microbe interactions and their effects on microbiome assembly and host function across taxonomic, molecular, genetic, and functional levels. Currently, bioinformatic tools identify and quantify bacterial isolates in SynComs by matching Amplicon Sequence Variants (ASVs) from microbiome datasets to *16S rRNA* amplicon sequences of SynCom members (Rognes et al., 2016; Zhang et al., 2021). This approach is suitable for low-complexity SynComs, where inoculated members are phylogenetically distant and easily distinguishable by their genomic and amplicon sequences. However, as SynComs increase in complexity and include more closely related members, accurate taxonomic resolution becomes more challenging, which in turn complicates functional analyses. Additionally, current bioinformatic tools do not account for intragenomic *16S rRNA* copy number variation, further skewing bacterial abundance estimates.

To address these limitations, we developed SynCom Fingerprinting (SyFi), a bioinformatics workflow that identifies sequence variants and copy number variation of target microbial genes or gene regions in bacterial genomes. SyFi uses these gene variants as a reference fingerprint for pseudoalignment-based quantification, inspired by methods originally developed for transcript isoform quantification in transcriptomics (Borozan et al., 2023; Patro et al., 2017). We demonstrate that SyFi improves accuracy in identifying, distinguishing, and quantifying SynCom members compared to current tools. Additionally, we compare microbiome compositions inferred by SyFi and other tools to metagenomic data, which serves as a gold standard. By improving the resolution of SynCom microbiome analyses, SyFi enables more complex SynCom studies, advancing our understanding of microbiome assembly and dynamics in plant-associated microbial communities.

## Methods

### SyFi workflow in brief

The SyFi workflow consists of three modules. In the first module (*SyFi main*), SyFi builds a fingerprint of each SynCom member utilizing the genome sequence and the genomic reads as input (Figure 1A-top & Appendix 1). Aside from the genome sequence and genomic reads, SyFi requires a complete target sequence from which a fingerprint will be built. In microbiome studies, for example, the conserved bacterial *16S rRNA* sequence is commonly used for metagenomic amplicon sequencing and could therefore be used as the target sequence. Here, we focus our analysis and description on the use of SyFi to obtain 16S rRNA fingerprints. However, other bacterial community markers, such as *recA* (Acinas et al., 2021), *rpoB* (Ogier et al., 2019), and *gyrB* (Barret et al., 2015) gene sequences can also serve as input for SyFi.

**Figure 1.**
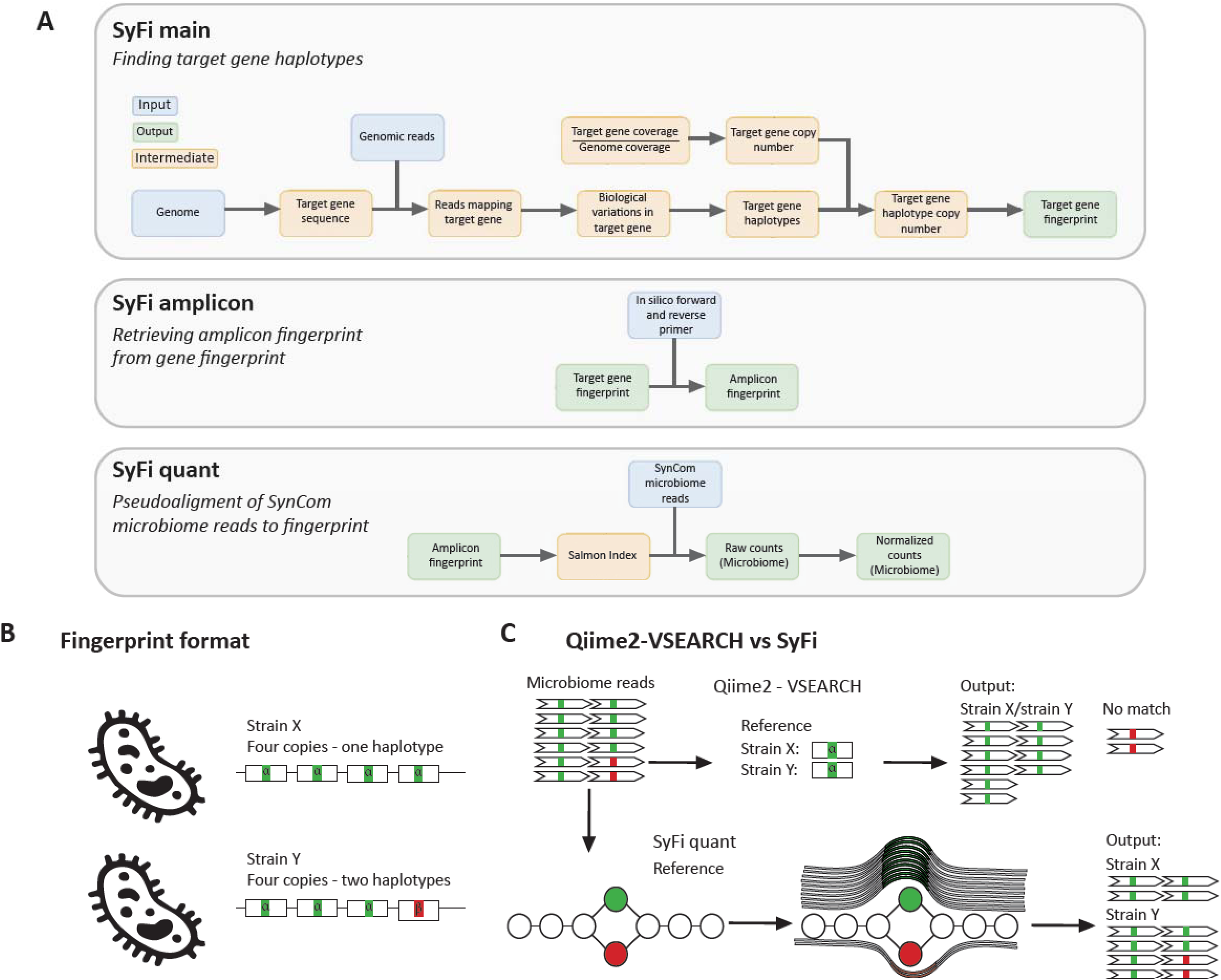
SyFi workflow and principle. A) The SyFi pipeline consists of three modules. *SyFi main* generates a unique fingerprint for each SynCom member based on a reference target sequence. This involves extracting the gene sequence from the genome and isolating corresponding reads from genomic data. These reads are used to identify biological variations and construct haplotype variants of the target gene sequence. The final fingerprint is constructed using these haplotypes and their copy numbers. *SyFi amplicon* utilizes *in silico* primers to extract the amplicon region from the target gene fingerprint. *SyFi quant* indexes the (amplicon) fingerprints, pseudoaligns SynCom microbiome reads to this reference, and normalizes raw counts based on the target gene’s copy number within the genomes. B) a schematic overview of fingerprint sequences, where two near-isogenic strains share an identical target gene and copy number, except for one haplotype variant in strain Y containing biological variation. C). Benchmark tools typically extract a single reference sequence per strain, usually the most commo haplotype. However, in near-isogenic strains, this approach may lead to indistinguishability. *SyFi quant* builds a De Bruijn graph k-mer index, pseudoaligns SynCom microbiome reads to the correct path within the graph and quantifies SynCom isolates. Small biological variations create splits in the De Bruijn graph, enabling higher resolution in distinguishing SynCom members.

Bacterial strains can harbor multiple copies of the *16S rRNA* gene (Sun et al., 2013), and these copies can have small biological variations like single nucleotide polymorphisms (SNPs), insertions, and deletions. SyFi is designed to identify all *16S rRNA* copies and biological variations in these *16S rRNA* sequences across a single bacterial genome. It aligns a representative sample *16S rRNA* sequence to the genome with the goal of retrieving the genomes’ *16S rRNA* sequence with the highest sequence identity score. Subsequently, SyFi uses raw genomic sequencing reads for read-based variant calling and phasing to determine within-target sequence biological variation and their co-occurrence. This information is used to assemble all unique *16S rRNA* sequences from the genome of interest, referred to as *16S rRNA* haplotypes. In the final step of the first module, SyFi determines the copy number of each *16S rRNA* haplotype by estimating the depth of coverage of each *16S rRNA* haplotype relative to the average genome-wide depth. Eventually, the sequences of all *16S rRNA* sequences from a given bacterial genome are concatenated into a fingerprint (Figure 1B). A detailed overview of the first module can be found in Appendix 1.

The second module of SyFi (*SyFi amplicon*) allows the user to extract an amplicon fingerprint from the gene fingerprint using *in silico* primers (Figure 1A-middle). Since the main SyFi module isn’t optimized for short target fragments, this module ensures the creation of fingerprints compatible with amplicon-sequenced SynCom datasets. In the third module (*SyFi quant*) (Figure 1A-bottom and 1C), the fingerprint is used as a reference for pseudoalignment of metagenomic amplicon sequencing reads using Salmon (version 1.4.0), to accurately identify SynCom members and quantify their abundance (Figure 1C) (Patro et al., 2017). Salmon indexes the fingerprints using k-mers, creating a data structure (De Bruijn graph) to efficiently match sequencing reads. Pseudoalignment then assigns reads to compatible fingerprints by identifying shared k-mers, without performing full sequence alignment. Thus, the *SyFi quant* module reports the number of reads that are pseudoaligned to each SynCom member’s fingerprint and, in the final step, normalizes these counts based on the SyFi-predicted *16S rRNA* copy number to estimate SynCom member abundances.

### 16S rRNA haplotype and copy numbers across bacterial taxa

To assess the prevalence of differential *16S rRNA* haplotypes in natural systems, we applied *SyFi main* to two large bacterial genomic datasets deriving from the human gut and plant root microbiomes (Forster et al., 2019; Selten et al., 2024b). *SyFi main* was run with default settings, using a, full-length *16S rRNA* seed sequence (1,544 bp) from a *Paenisporosarcina* strain as the target (Supplementary data S1), which is approximately the proposed length of the *16S rRNA* sequence of 1,550 bp (Clarridge, 2004). To maximize the recovery of complete *16S rRNA* genes across bacterial isolates, we allowed a maximum length deviation of 300 base pairs accounting for natural variation in *16S rRNA gene* length.

### SyFi performance on a SynCom-inoculated root microbiome dataset

To assess the performance of SyFi with relation to SynCom member quantification in microbiome datasets, we made use of a complex SynCom dataset that included the bacterial genomes from the same Selten et al. (2024b) bacterial genome dataset. This dataset consists of 44 samples in which three different plant hosts were inoculated with a SynCom consisting of 447 different bacteria. The isolation, whole genome sequencing, genome assembly of the bacteria, and the experimental set-up of the SynCom reconstitution experiment is described in (Selten et al., 2024a).

To quantify the abundance of the 447 SynCom members in these 44 samples, we generated *16S rRNA* V3-V4 and V5-V7 amplicon sequencing data (Table S1) and performed untargeted shotgun metagenome sequencing (Selten et al., 2024a) (Appendix 2). Pseudoalignment of amplicon reads to SyFi-generated fingerprints is most effective when the reference sequences correspond to the exact amplicon region covered by the reads (Bray et al., 2016) (Table S2). However, for accurate fingerprint generation, SyFi benefits from using the full-length target gene, as it requires a sufficient number of genomic reads mapping to the target gene to identify statistically supported biological variation. Therefore, we trimmed the already generated full-length *16S rRNA* fingerprints to V3-V4 and V5-V7 *16S rRNA* fingerprints using *SyFi amplicon* that utilizes the Qiime2’s read extraction algorithm (version 2022.8) (Bolyen et al., 2019). To create *16S rRNA* fingerprints for the V3-V4 and V5-V7 regions, we used the following primers in *SyFi amplicon*: forward primer for V3-V4: CCTACGGGNGGCWGCAG, reverse primer for V3-V4: GACTACHVGGGTATCTAATCC, forward primer for V5-V7: AACMGGATTAGATACCCKG, and reverse primer for V5-V7: ACGTCATCCCCACCTTCC (Beckers et al., 2016).

### Quantification of SynCom members using the SyFi fingerprints

To quantify the SynCom members, we evaluated pseudoalignment of the amplicon reads to the V3-V4 and V5-V7 *16S rRNA* fingerprints. The raw reads of the V3-V4 and V5-V7 *16S rRNA* amplicon datasets were filtered, denoised, and low-quality bases were trimmed at position 20 and 240 at the 3’ end and 5’ end respectively for both forward and reverse reads using DADA2 (DADA2 version 1.26.0) (Callahan et al., 2016). The *16S rRNA* V3-V4 and V5-V7 amplicon data was pseudoaligned to the *16S rRNA* V3-V4 and V5-V7 fingerprints with Salmon (version 1.4.0) with default options except for a minimum mapping read identity score of 0.95 (Patro et al., 2017). The number of pseudoaligned reads was subsequently normalized for *16S rRNA* copy number that was calculated in the first module.

### Quantification of SynCom members using Qiime2 and VSEARCH

To compare SyFi’s accuracy in identifying, distinguishing, and quantifying SynCom members, we also preprocessed the *16S rRNA* V3-V4 and V5-V7 datasets with Qiime2 and DADA2, and quantified the SynCom members with VSEARCH. This is the current standard for profiling microbiomes in SynCom reconstitution experiments and is therefore used here as a benchmark. The amplicon reads were imported into Qiime2 and denoised by DADA2 as before (Qiime2 version 2022.11) (Bolyen et al., 2019). The ASV sequences were matched with the reference V3-V4 or V5-V7 *16S rRNA* amplicon sequences using the Qiime2-VSEARCH plugin (Rognes et al., 2016). These references were retrieved from the *16S rRNA* sequence from SyFi’s first step (Figure 1; BLAST alignment (v2.13.0)) with Qiime2 (version 2022.8) using the previously described V3-V4 and V5-V7 *16S rRNA* primers that were used in *SyFi amplicon*.

### Data analysis

To investigate SyFi’s ability to distinguish isolates based on *16S rRNA*, V3-V4, or V5-V7 fingerprints, we clustered the fingerprints at 100% sequence similarity with VSEARCH (VSEARCH version 1.2.13) (Rognes et al., 2016). The established clusters of isolates were subsequently used to compare the bacterial abundances in the fingerprint dataset with the abundances in the metagenome-sequenced dataset. The abundances of isolates of the same cluster were summed in both the fingerprint and metagenome dataset to ascertain a fair comparison of identical isolates. Afterwards, the abundances were normalized to relative counts and isolates that were absent in either dataset were removed in both the fingerprint and metagenome dataset.

Pearson correlations were calculated between the fingerprint and metagenome-sequenced datasets to assess SyFi’s accuracy in quantifying isolates. To assess whether SyFi correctly assigned presence or absence to an isolate compared to the metagenome dataset, a presence/absence threshold was implemented in both datasets. Strains with a minimum relative abundance of 0.5% were assigned as being present.

All data analysis steps described above were performed identically for the VSEARCH analyses of the SynCom dataset as well as the Salmon run on the original amplicon sequences.

## Results

### The principle of SyFi

SyFi is inspired and partially built upon the principle of pseudoalignment-based quantification methods used in transcriptomics. This approach involves creating a kmer-based reference for near-isogenic gene isoforms, which can be distinguished through pseudoalignment. In this method, reads are matched to the correct isoform by aligning with the appropriate k-mer paths in a De Bruijn graph (Figure 1C). Minor biological variations in a near-isogenic bacterial strain’s gene reference cause splits in these k-mer paths, enabling the accurate identification of the correct isoform or SynCom member, even when they are highly similar.

SyFi consists of three modules. The first module*, SyFi main* (Figure 1A-top and Appendix 1), constructs a genomic fingerprint for each micro-organism to be quantified later. *SyFi main* uses the genome and genomic reads of bacterial strains to identify variants of a target sequence and builds a fingerprint consisting of a concatenation of all detected gene variants and/or copies, here illustrated for the *16s rRNA gene*. Target sequence copies can be identical, but can also differ in their sequence, frequently referred to as haplotypes, due to single nucleotide polymorphisms (SNPs), insertions, or deletions (Figure 1B). The second module, *SyFi amplicon* (Figure 1A-middle), extracts regions from the fingerprints, essential for amplicon sequence analysis. Such analysis usually focuses on a smaller region of the target gene, for example a targeted variable region (or regions) of the *16S rRNA* gene, that can be captured by short-read sequencing. Lastly, the third module, SyFi quant, quantifies microbial abundance by leveraging biological variation between amplicon sequences (Figure 1A-bottom and 1C). This is achieved by pseudoalignment-based quantification, implemented in Salmon (Patro et al., 2017). By distinguishing highly similar fingerprints from closely related microbes, this approach enables more accurate quantification, analogous to the analysis of alternative splicing in transcriptomics (Fu et al., 2019) (Figure 1C).

### SyFi distinguishes more bacteria

To assess whether SyFi’s ability to distinguish bacteria more effectively than standard methods, we evaluated *SyFi main*’s performance on two large bacterial genomic datasets (Forster et al., 2019; Selten et al., 2024b) and *SyFi quant* ’s performance on a complex SynCom dataset (Selten et al., 2024a). SyFi was applied to genome sequences of 710 human gut- and 447 *Arabidopsis thaliana* (hereafter Arabidopsis) root-derived bacterial isolates, successfully retrieving *16S rRNA* fingerprints for 458 and 382 isolates, respectively (Table 1). The isolates for which SyFi was unable to construct a fingerprint either lacked the *16S rRNA* gene or had an incomplete gene sequence.

**Table 1.**
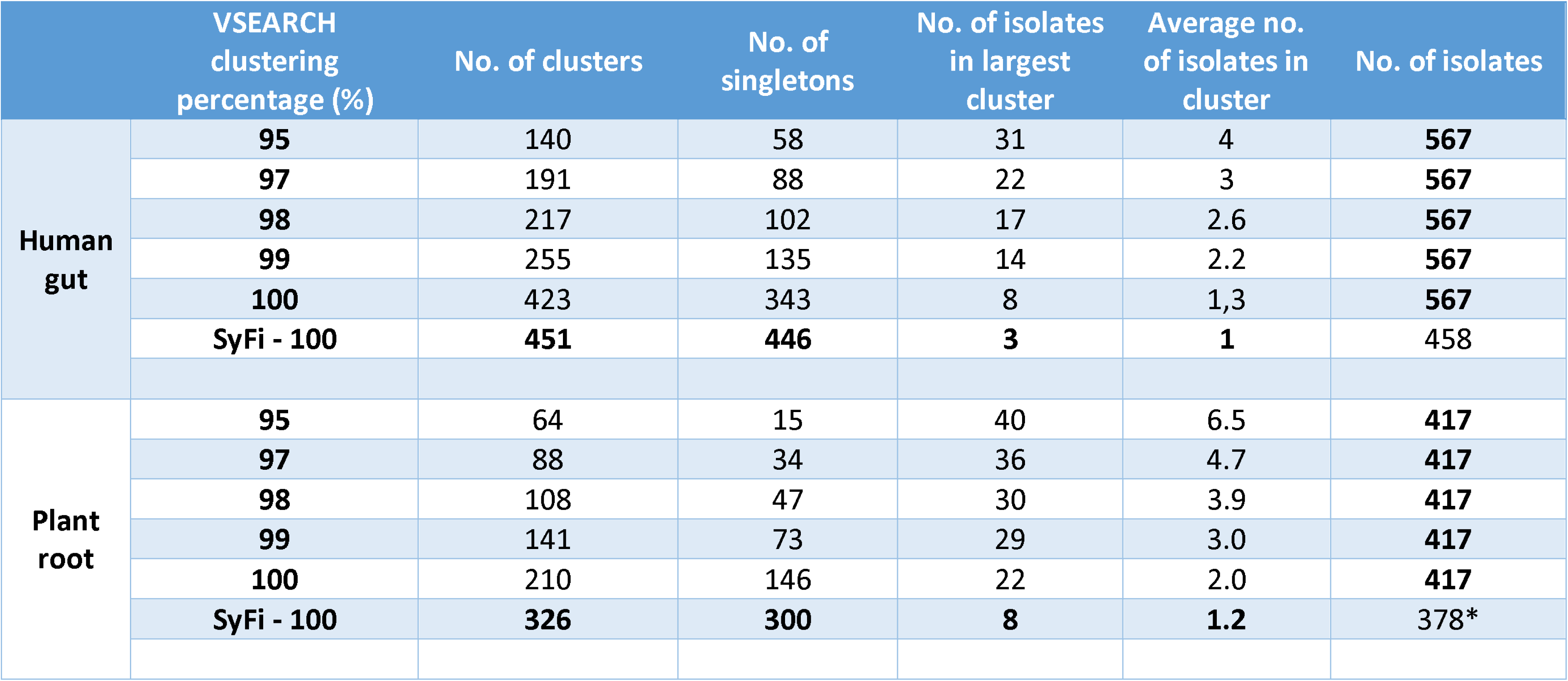
Comparison of SyFi-generated 16S rRNA fingerprints and directly retrieved 16S rRNA sequences in distinguishing bacterial isolates. The first column indicates whether VSEARCH was applied to the original 16S rRNA sequences extracted directly from the bacterial genomes, clustered at varying identity thresholds (95%, 97%, 98%, 99% and 100%), or to the SyFi- generated fingerprints clustered at 100% sequence identity (SyFi - 100). *Note: Although 382 16S rRNA fingerprints were obtained, four sequences were excluded due to exceeding VESEARCH’s maximum sequence length threshold.

|  | VSEARCH<br>clustering<br>percentage (%) | No. of clusters | No. of<br>singletons | No. of isolates<br>in largest<br>cluster | Average no.<br>of isolates in<br>cluster | No. of isolates |
| --- | --- | --- | --- | --- | --- | --- |
| Human<br>gut | 95 | 140 | 58 | 31 | 4 | 567 |
|  | 97 | 191 | 88 | 22 | 3 | 567 |
|  | 98 | 217 | 102 | 17 | 2.6 | 567 |
|  | 99 | 255 | 135 | 14 | 2.2 | 567 |
|  | 100 | 423 | 343 | 8 | 1,3 | 567 |
|  | SyFi - 100 | 451 | 446 | 3 | 1 | 458 |
| Plant<br>root | 95 | 64 | 15 | 40 | 6.5 | 417 |
|  | 97 | 88 | 34 | 36 | 4.7 | 417 |
|  | 98 | 108 | 47 | 30 | 3.9 | 417 |
|  | 99 | 141 | 73 | 29 | 3.0 | 417 |
|  | 100 | 210 | 146 | 22 | 2.0 | 417 |
|  | SyFi - 100 | 326 | 300 | 8 | 1.2 | 378* |

To determine whether genome assembly fragmentation contributed to the missing *16S rRNA* sequences, we compared genome completeness, coverage, and L50—three indicators of genome assembly integrity—between isolates with and without SyFi-generated 16S rRNA fingerprints for the plant root bacterial genome dataset (Figure S1). Isolates with SyFi-identified fingerprints had significantly higher genome completeness and coverage, while those without fingerprints exhibited greater genome fragmentation (low L50) and a higher number of scaffolds. This suggests that incomplete or fragmented genomes indeed reduce SyFi’s ability to retrieve *16S rRNA* fingerprints.

We next assessed the discriminative power of SyFi-generated *16S rRNA* fingerprints using VSEARCH clustering at 100% sequence identity (Table 1). This analysis identified 451 and 326 non-redundant *16S rRNA* fingerprints for the human gut and plant root datasets, respectively, with 446 and 300 being unique. In comparison, direct alignment-based clustering retrieved a higher number of *16S rRNA* sequences – 567 for the human gut and 417 for the plant root dataset – but these included partial and fragmented sequences. In spite of that difference, we found only 423 and 210 non-redundant sequences for the human gut and plant root datasets, with just 343 and 146 being unique, respectively. Overall, SyFi improved isolate resolution across both datasets distinguishing 98.5% (451/458) of human gut isolates and 86.2% (326/378) of plant root isolates. This was a substantial improvement over standard methods, which identified only 74.6% (423/567) and 47.0% (210/417) of isolates, respectively. Furthermore, SyFi outperformed current tools not only at the full-length *16S rRNA* level but also within the V3-V4 and V5-V7 amplicon regions (Table S3).

### SyFi identifies *a* similar 16S rRNA copy number distribution across the bacterial kingdom

For both the human gut and the plant root, SyFi identified most bacterial genomes as having two to five *16S rRNA* copies (Figure 2A and 3A), with the majority containing at least two distinct haplotypes (Figure 2B and 3B), a feat especially present in the human gut microbiome. These findings align with previous analyses of over 2,000 bacterial genomes (Sun et al., 2013; Větrovský & Baldrian, 2013). Standard direct alignment methods, by contrast, typically detect only a single *16S rRNA* haplotype per genome (Figure S2), illustrating how conventional approaches often overlook intragenomic variation.

**Figure 2.**
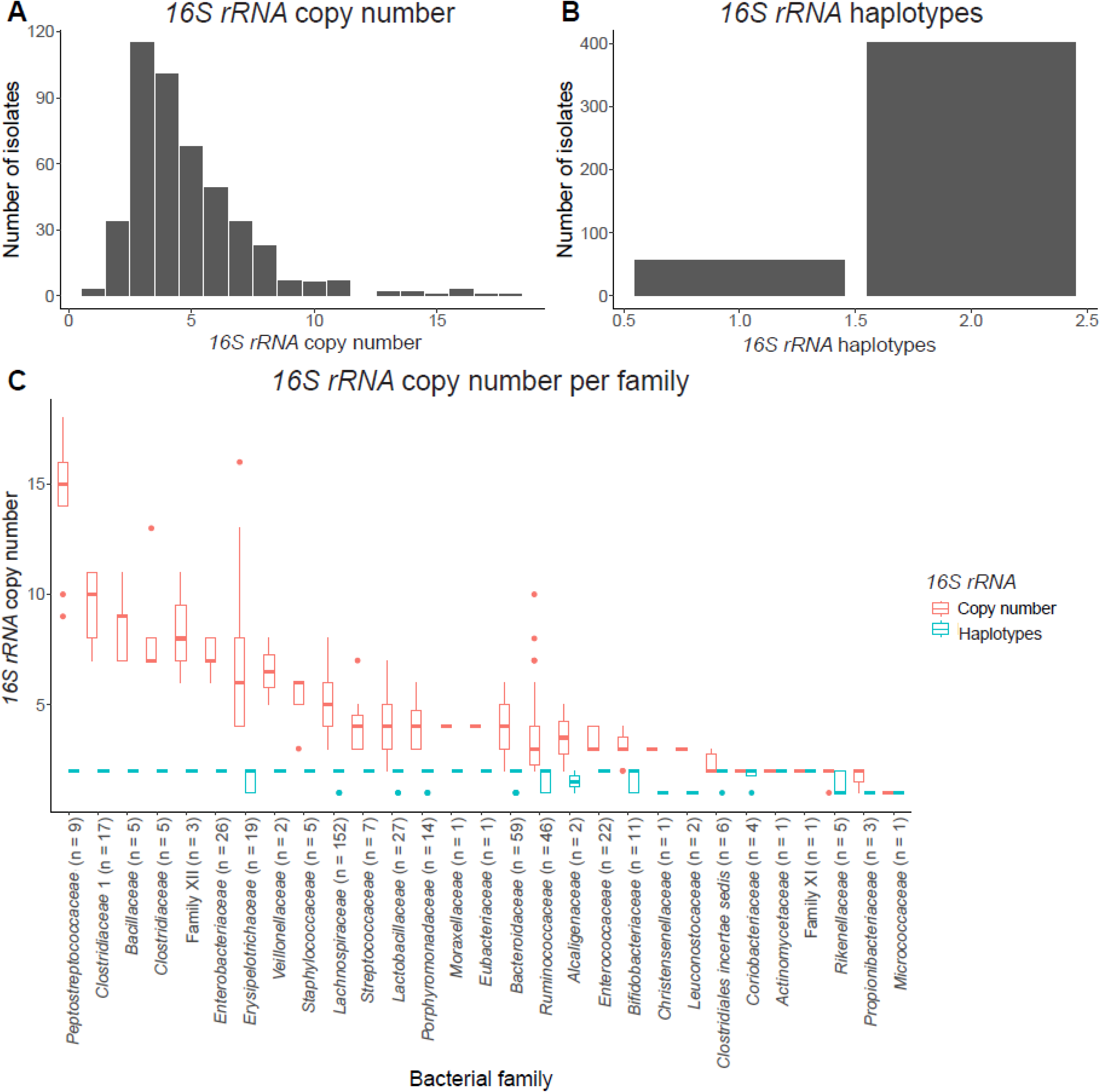
*16S rRNA* copy number and haplotype distribution among 458 human gut bacterial isolates. (A) *16S rRNA* copy number. (B) *16S rRNA* haplotype distribution. (C) Distribution of *16S rRNA* copy number (red) and the number of *16S rRNA* haplotypes (turquoise) within diverse bacterial families. Families are ordered by the average number of *16S rRNA* copies in descending order. The number in brackets indicates the number of isolates in the respective family.

**Figure 3.**
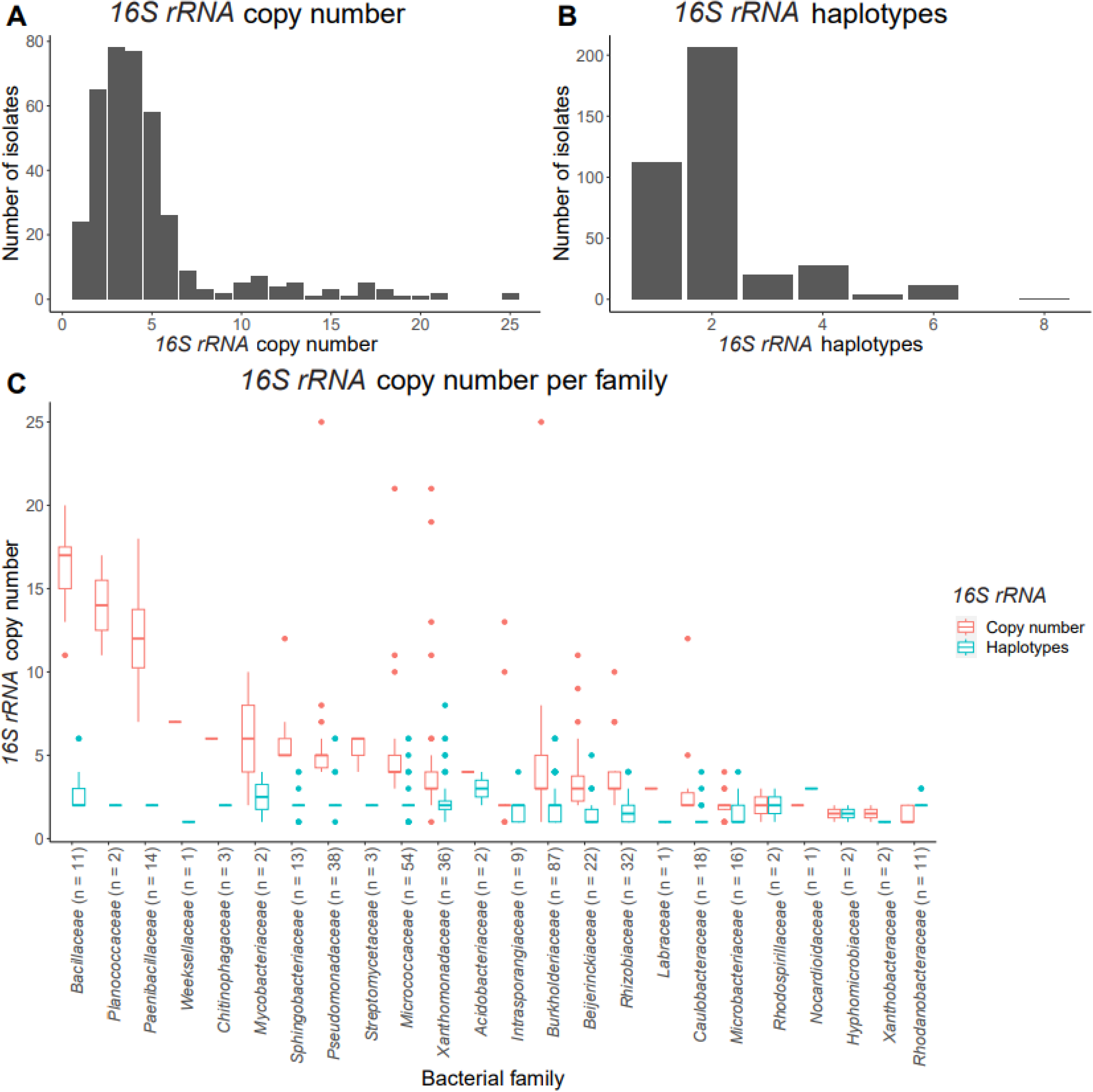
*16S rRNA* copy number and haplotype distribution among 382 bacterial isolates. (A) *16S rRNA* copy number. (B) *16S rRNA* haplotype distribution. (C) Distribution of *16S rRNA* copy number (red) and the number of *16S rRNA* haplotypes (turquoise) within diverse bacterial families. Families are ordered by the average number of *16S rRNA* copies in descendin order. The number in brackets indicates the number of isolates in the respective family.

To validate these observations with recent data, we analyzed 3,615 complete bacterial genomes retrieved from NCBI (April 15, 2024), spanning the entire bacterial kingdom (Figures S3A and B). Consistent with prior studies, we found that most genomes contain two to five 16S rRNA copies (90.7%) but only one detected haplotype (Sun et al., 2013; Větrovský & Baldrian, 2013). Additionally, across all datasets, the bacterial families *Bacillaceae*, *Planococcaceae* and *Paenibacillaceae* had the highest *16S rRNA* copy numbers (Figures 2C, 3C, S3C, and S3D), reinforcing SyFi’s accuracy in estimating *16S rRNA* copy number.

The prevalence of only one detected haplotype in complete bacterial genomes is surprising. Bacterial genome assembly often relies on Illumina short-read sequencing combined with PacBio or Nanopore long-read sequencing (Sereika et al., 2022). However, these methods may interpret true sequence variation as sequencing errors and erroneously correct them, potentially explaining the limited haplotype diversity observed. Although genome-polishing tools are improving (Luan et al., 2024), many complete genome assemblies remain unpolished. To investigate whether there is an underestimation of the number of *16S rRNA* haplotypes in the complete genomes, we analyzed the 16S rRNA gene(s) in ten Bacillales strains with complete genomes and for which raw Illumina short reads were available. BLASTn alignment of the *16S rRNA* gene (Supplementary data S1) to these ten genomes, indeed, yielded a single *16S rRNA* haplotype for nine out of ten strains. However, alignment of the corresponding short read sequences by SyFi yielded two distinct haplotypes in all ten cases (Table S4) with clear biological variations at specific positions in the *16S rRNA* sequence (Figure S4). This highlights an important limitation in genome assembly, where complete genomes potentially lack biological variation, whereas SyFi can recover this variation, providing a more accurate overview of sequence diversity.

While complete genomes may underestimate *16S rRNA* haplotype diversity, fragmented or draft genome assemblies are more susceptible to contamination, which can affect the construction of *16S rRNA* fingerprints. A detailed analysis of contamination, genomic heterogeneity, and GC content indicates how SyFi may slightly overestimate *16S rRNA* copy numbers with moderate to high contamination and/or heterogeneity levels (Appendix 3). Even less favorable is how high GC contents may exponentially overestimate *16S rRNA* copy numbers and future implementations of SyFi may be adjusted for these (Appendix 3).

### SyFi outperforms VSEARCH in identifying and quantifying bacterial isolates

To assess SyFi’s accuracy in profiling SynCom-inoculated microbiome samples, we analyzed a complex SynCom dataset comprising 447 bacterial isolates inoculated on plant roots (Selten et al., 2024a). We sequenced the V3-V4 and V5-V7 *16S rRNA* amplicons from the root microbiome samples and pseudoaligned the amplicon reads to SyFi-generated *16S rRNA* fingerprints using *SyFi quant*. First, we optimized pseudoalignment accuracy by testing various minimum sequence similarity thresholds in Salmon while maximizing the number of pseudoaligned reads (Figures S5 and S6). We then compared SyFi-based quantifications to metagenome sequencing—considered the gold standard for validating amplicon sequencing datasets (Brumfield et al., 2020; Laudadio et al., 2018; Rausch et al., 2019). SyFi achieved 93.6% and 81.0% accuracy in quantifying isolate abundances in the V3-V4 and V5-V7 datasets, respectively (Figures 4, S7, and Table S5). Furthermore, SyFi correctly assessed isolate presence/absence with 92.8% and 92.1% accuracy for the V3-V4 and V5-V7 datasets, respectively (Table S5).

**Figure 4.**
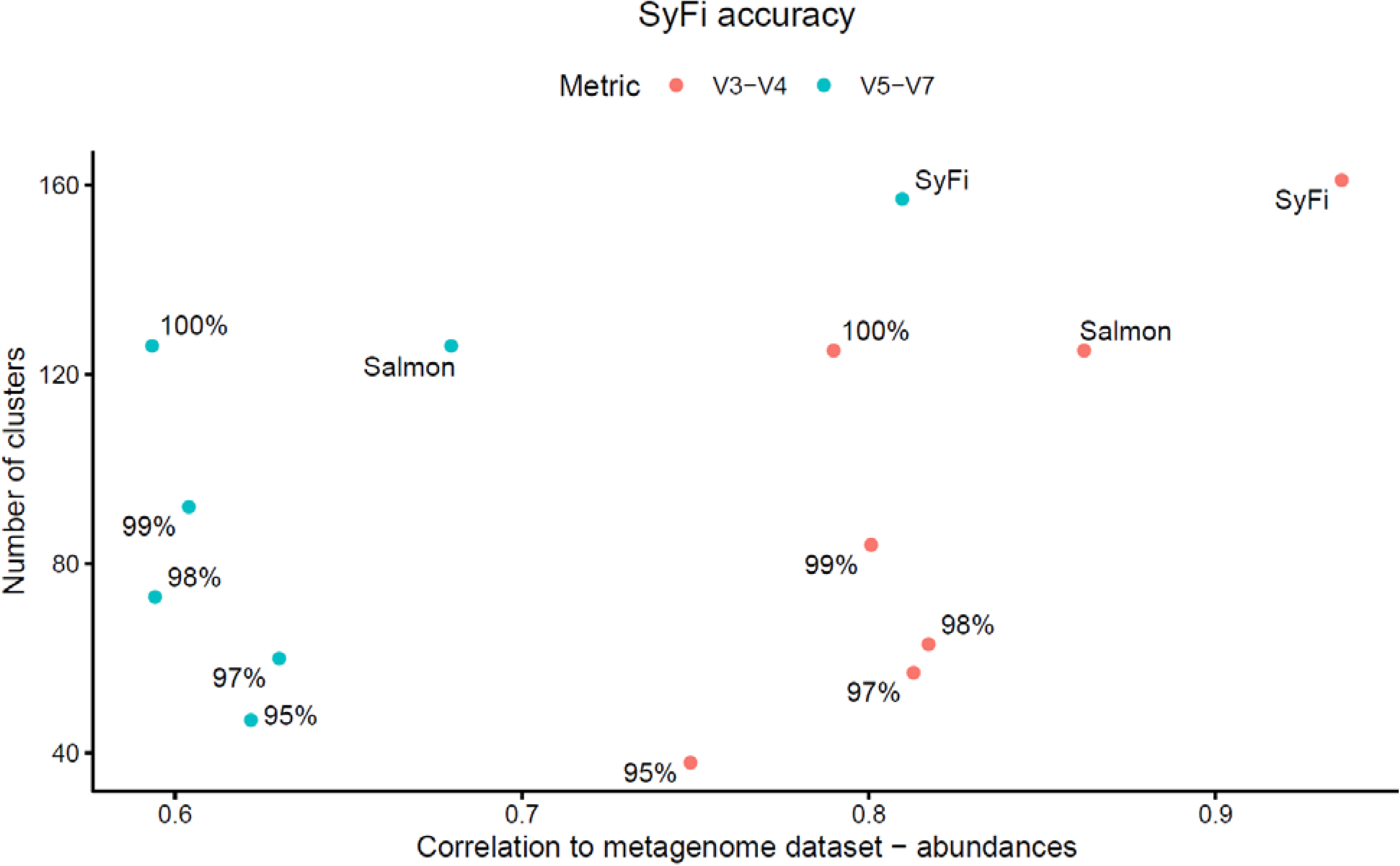
Accuracy of SyFi in quantifying bacterial isolate abundances. The coordinates indicate the sequence identity percentage at which the amplicons were clustered by VSEARCH, and if the coordinate is from direct pseudoalignment (Salmon) or SyFi. The y-axis shows the maximum number of distinguishable isolates in the dataset based on sequence identity, while th x-axis displays the Pearson correlation value between the dataset and the metagenome-based quantification, reflectin accuracy in estimating isolate abundances.

We then compared SyFi’s performance to the Qiime2-DADA2 pipeline with VSEARCH-based matching. SyFi outperformed the Qiime2-VSEARCH method (100% sequence identity threshold) in quantification accuracy for both the V3-V4 (93.6% vs. 79.0%) and V5-V7 (81.0% vs. 59.3%) datasets (Figure 4). SyFi also marginally improved presence/absence calling in the V3-V4 dataset (92.8% vs. 90.8%) but was slightl outperformed for presence/absence calling in the V5-V7 dataset (92.1% vs. 94.4%) (Table S5). Nevertheless, SyFi identified more isolates overall (157 vs. 126), suggesting that while Qiime2-VSEARCH may more accurately assign amplicons to clusters of isolates, SyFi provides finer resolution within clusters by leveraging the V5-V7 fingerprint.

To determine the contribution of pseudoalignment-based quantification to the improved performance of SyFi, we pseudoaligned V3-V4 and V5-V7 amplicon reads to SynCom reference 16S rRNA sequences alone (Figure 4; “Salmon coordinates”). Across both regions, SyFi consistently improved pseudoalignment accuracy (quantification: V3-V4, 93.6% vs. 86.2%; V5-V7, 81.0% vs. 68.0%; identification: V3-V4, 92.8% vs. 92.0%; V5-V7, 92.1% vs. 91.0%) (Figure 4 and Table S5).

Overall, SyFi more accurately identifies and quantifies bacterial isolates than VSEARCH or pseudoalignment alone in both V3-V4 and V5-V7 amplicon datasets (Figure 4 and Table S5). Additionally, SyFi quant yields similar or higher pseudoaligned read counts compared to Qiime2-VSEARCH (Figure S6).

Click or tap here to enter text.

## Discussion

As SynCom complexity increases, accurately quantifying bacterial strains using marker gene-based amplicon sequencing becomes more challenging. To address this, we developed SyFi (SynCom Fingerprinting). SyFi is designed to identify intragenomic sequence variation in bacterial genomes for a given marker gene and use the resulting sequence variants to construct genomic fingerprints that in turn can be used to quantify the abundance of microbes in a complex community at heightened taxonomic resolution. Additionally, we show that pseudoalignment of amplicon reads to SyFi-generated fingerprints outperforms classical methods in identifying, distinguishing, and quantifying bacterial isolates. Beyond the dissection of complex SynCom datasets, SyFi can be applied to investigate sequence variability in any multi-copy gene.

A computational screen of *16S rRNA* copy number and haplotype distribution across bacterial genomes reveals that genome assemblies often miss small variations in the *16S rRNA* sequence. In this context, SyFi can capture these sequence variations and more accurately determine which *16S rRNA* haplotypes are present compared to retrieving *16S rRNA* sequences directly from genomes using target alignment. The *16S rRNA* copy number distribution generated by SyFi aligns with previous studies (Sun et al., 2013; Větrovský & Baldrian, 2013) and indicates for both human gut and plant root microbiomes that bacteria often harbor more than one *16S rRNA* haplotype sequence (Figures 2 and 3). Surprisingly, we discovered few *16S rRNA* haplotypes in complete genomes when retrieving them directly (Sun *et al*., 2013), which suggests SyFi might be identifying PCR errors as biological variations in *16S rRNA* sequences. However, when ran on ten closed Bacillales genomes with a single *16S rRNA* haplotype, SyFi was able to detect multiple sequence variants. This highlights a major limitation in the current closed bacterial genomes in databases. The usage of long reads to concatenate and polish short-read-assembled genomes leads to the loss of sequence variants across multi-copy marker genes. Despite this, advancements in genome- polishing tools are improving the accuracy of complete bacterial genomes, e.g. by assembling genomes with long reads instead of short reads and using the short reads to infer biological variations (Luan *et al*., 2024). This effort makes closed genomes increasingly suitable for complex SynCom studies.

SyFi’s ability to extract accurate *16S rRNA* haplotype profiles from a bacterial genome depends on the quality of the genome assembly and the availability, coverage, and accuracy of genomic reads (Appendix 1 and 3). Contaminating sequences in the genome assembly or genomic reads may lead to an overestimation of *16S rRNA* copy numbers or the presence of highly dissimilar intragenomic *16S rRNA* sequences, which might seem unlikely to co-occur in a single genome. Previous studies have shown that the vast majority of *16S rRNA* sequences conform to >95% intra-species sequence identity (Jain et al., 2018), suggesting that *16S rRNA* haplotypes with <95% sequence identity within a genome are improbable. In rare cases, highly divergent *16S rRNA* sequences may arise due to events such as horizontal gene transfer or the uptake of DNA fragments from the environment (Kitahara et al., 2012; Miyazaki & Tomariguchi, 2019). Beyond data quality, *16S rRNA* fingerprints may also be subject to copy number overestimation due to the higher GC content of *16S rRNA* sequences relative to the rest of the genome (Appendix 3). Whether this affects microbiome composition accuracy remains uncertain, as the GC content-derived biases may be present in both amplicon and metagenomic datasets (Browne et al., 2020). Consequently, the GC content bias in genomic data may be counterbalanced by a similar bias in metagenomic data. Future versions of SyFi may incorporate an optional *16S rRNA* copy number normalization step based on differential GC content.

Additional features for future SyFi versions may include modifications to the input target sequence and refinements to the haplotype calling step (Martin et al., 2016). We have not yet assessed whether different *16S rRNA* target sequences significantly alter resulting *16S rRNA* fingerprints. Targeting a shorter or less diverse *16S rRNA* sequence may yield shorter fingerprints and/or lower resolution in distinguishability, potentially influencing microbiome composition as processed through *SyFi quant*. To better understand the impact of target sequence selection on *16S rRNA* fingerprints and microbiome composition, future versions of SyFi could incorporate a phylogeny-dependent marker gene database, allowing target sequence selection based on isolate phylogeny. A set of complete *16S rRNA* sequences spanning the bacterial kingdom will allow the retrieval of the most complete *16S rRNA* sequences for every tested SynCom member. Additionally, SyFi currently employs the “WhatsHap phase” algorithm for *16S rRNA* read phasing, which assumes a diploid model with only two haplotype outcomes. This assumption may lead to an underestimation of the true haplotype diversity within bacterial genomes, especially in cases of extensive biological variation. Despite this limitation, SyFi outperforms benchmark microbiome profiling methods. However, future versions could further improve enhance accuracy by enabling phasing beyond two haplotypes per sequence, e.g. by using the polyploidy phasing algorithm (Schrinner et al., 2020), (providing a more precise representation of intragenomic haplotype diversity.

Over the past decade, increasing evidence has highlighted substantial variability in marker genes, such as the *16S rRNA* gene, in terms of copy number and sequence identity. Although rarely accounted for in microbiome profiling, this variability impacts the estimation of bacterial presence and abundance in natural microbiomes. High *16S rRNA* copy numbers can lead to an overestimation of bacterial strain abundance in microbiome datasets (Klappenbach, 2001; Louca et al., 2018; Pei et al., 2010; Sun et al., 2013; Větrovský & Baldrian, 2013), while multiple intragenomic *16S rRNA* variants can result in an overestimation of bacterial species diversity (Schloss, 2021). Therefore, considering marker gene variability in microbiome studies is crucial for improving dataset accuracy. In SynCom studies, the primary objective is to accurately identify and quantify bacteria within the dataset. While SyFi does not perfectly distinguish all bacterial isolates in amplicon sequencing datasets, it represents a significant advancement in microbiome profiling by improving the accuracy of bacterial identification and quantification.

## Acknowledgments and funding

The authors want to thank members of the Plant-Microbe Interactions lab (Utrecht, The Netherlands) and the Molecular Biology and Genetics lab (Aarhus, Denmark) for helpful discussions and to Juan J. Sanchez-Gil for brainstorming on the methodology and providing critical feedback on the manuscript. This study was supported by the Novo Nordisk Foundation Grant no. NNF19SA0059362. We acknowledge Novogene for providing sequencing service and data.

## Competing interests

The authors declare no competing interests.

## Author contributions

G.S. and R.d.J. conceptualized the methodology for SyFi. G.S. and A.G.-R. made the graphical visualization of the methodology. G.S. and A.G.-R developed the code, incorporating suggestions and insights from R.d.J. F.L. generated the data used for SyFi’s validation, which G.S. investigated and analyzed. S.R. and R.d.J. oversaw project administration and supervision. The manuscript was written by G.S. and R.d.J., with all the authors contributing to discussions and content development.

## Supplementary Data

Scripts and data generated to produce the figures in the manuscript or appendices can be found on https://github.com/adriangeerre/SyFi.

## Supplementary figures

**Figure S1.**
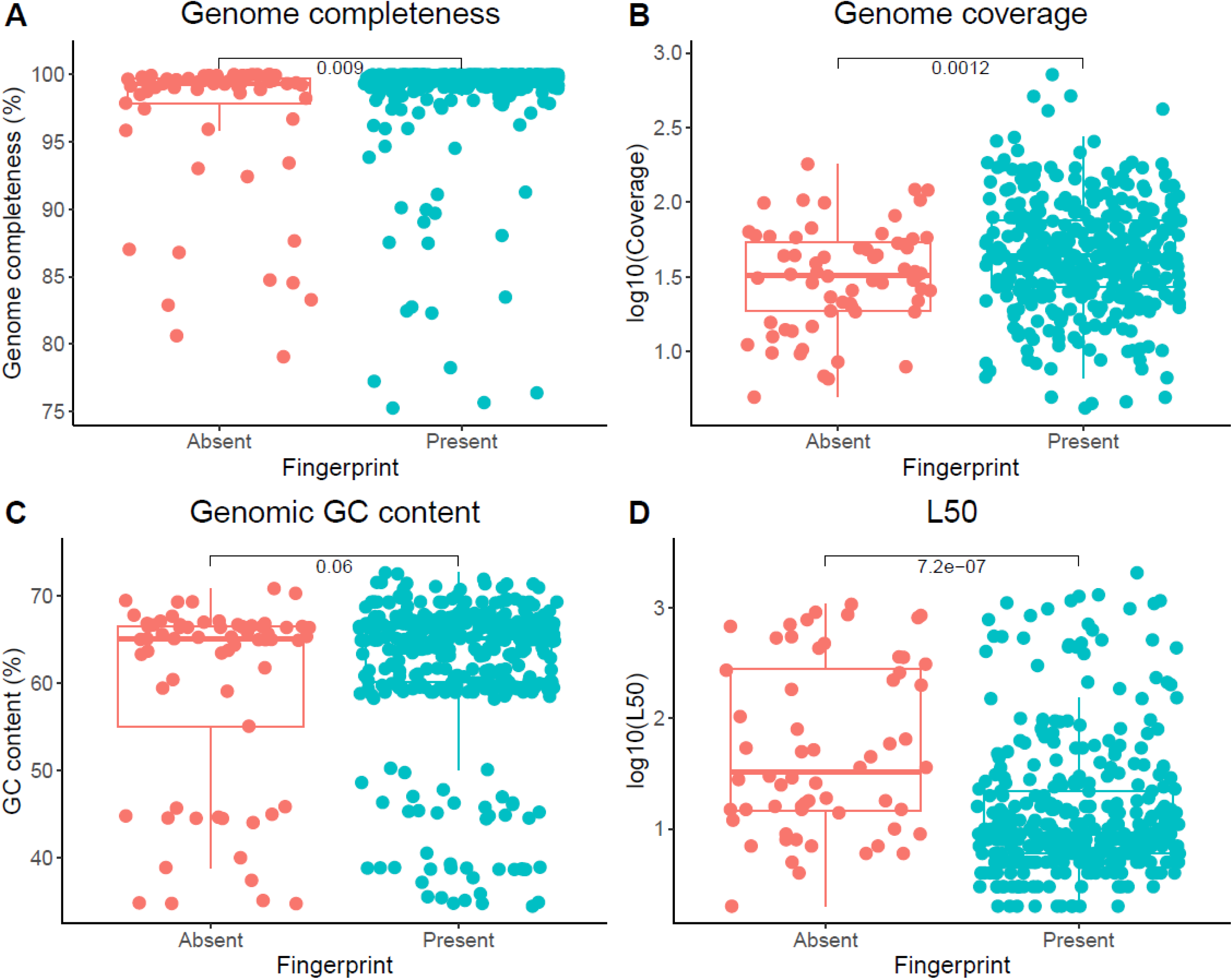
Effect of genome and genome assembly features on SyFi’s ability to build a *16S rRNA* fingerprint. Genome completeness (A), coverage (B), GC content (C) and L50 (D) of the genomes for which SyFi was able to build a *16S rRN A* fingerprint (Present) or not (Absent). Statistical differences are tested with a pairwise t-test.

**Figure S2.**
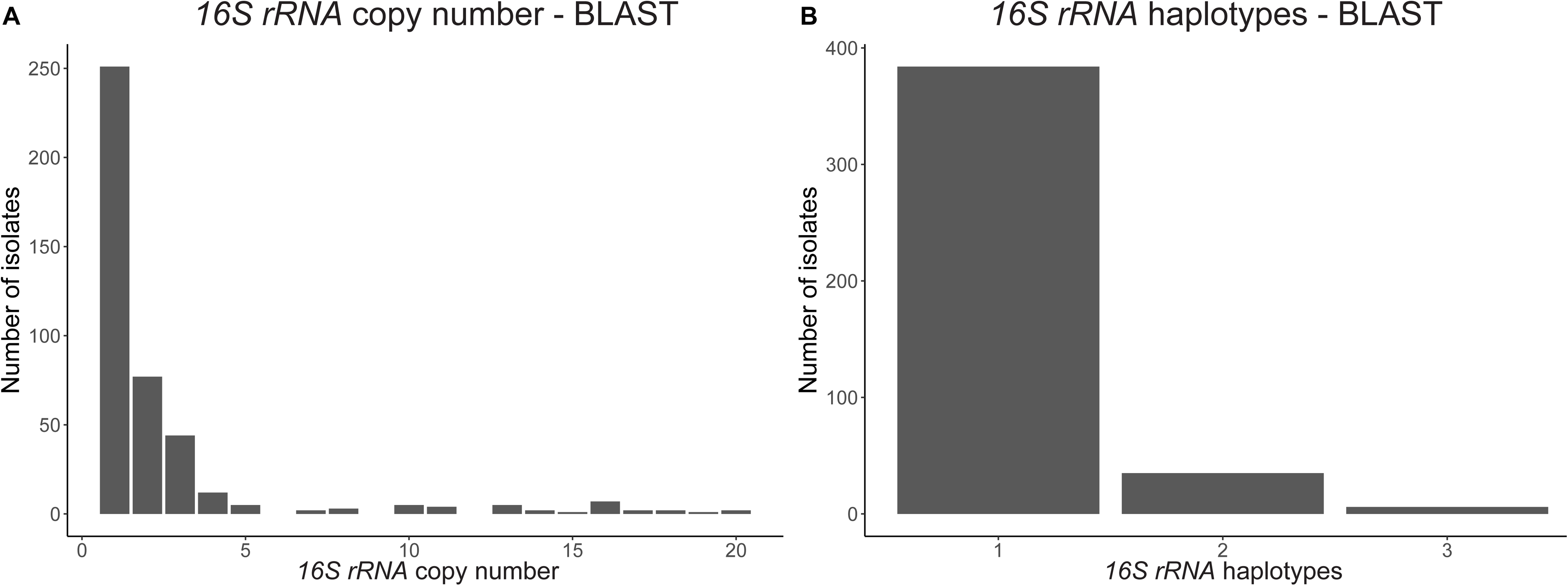
*16S rRNA* copy number and haplotype distribution among the 447 plant root isolates using BLAST to retrieve th *16S rRNA* sequences. The *16S rRNA* copy number (A) and number of *16S rRNA* haplotypes (B) when using a direct alignment of the target gene to the bacterial genomes.

**Figure S3.**
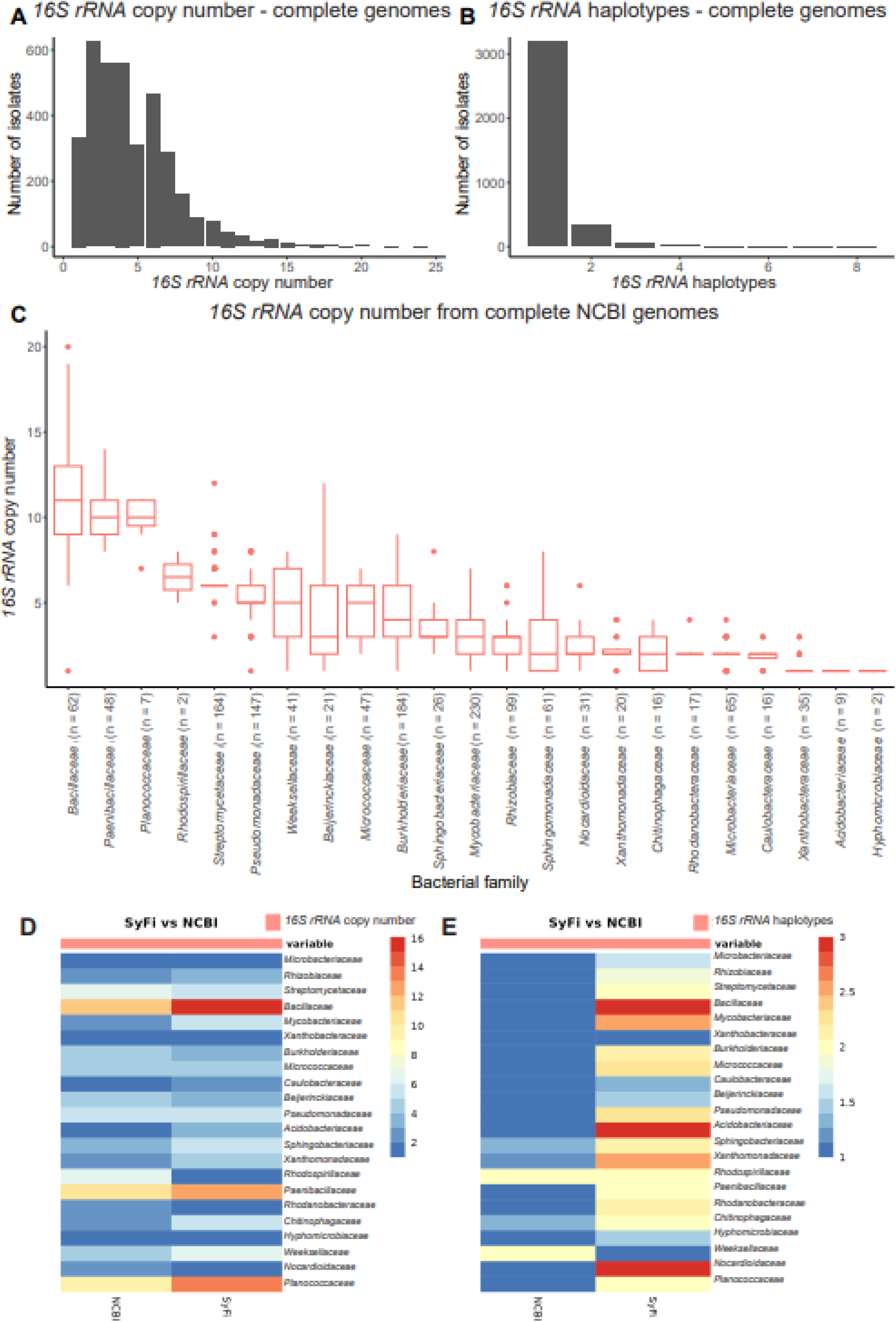
The *16S rRNA* copy number and haplotypes in complete NCBI genomes. The *16S rRNA* copy number (A) and 1*6S rRNA* haplotypes (B) in 3,615 complete NCBI genomes. The *16S rRNA* copy number per family (C) in the complete NCBI genomes subsetted for families that are present among the 447 isolates (Figure 3C). Comparison of *16S rRNA* copy number (D) and *16S rRNA* haplotypes (E) between the 447 plant root isolates and the 3,615 complete NCBI genomes.

**Figure S4.**
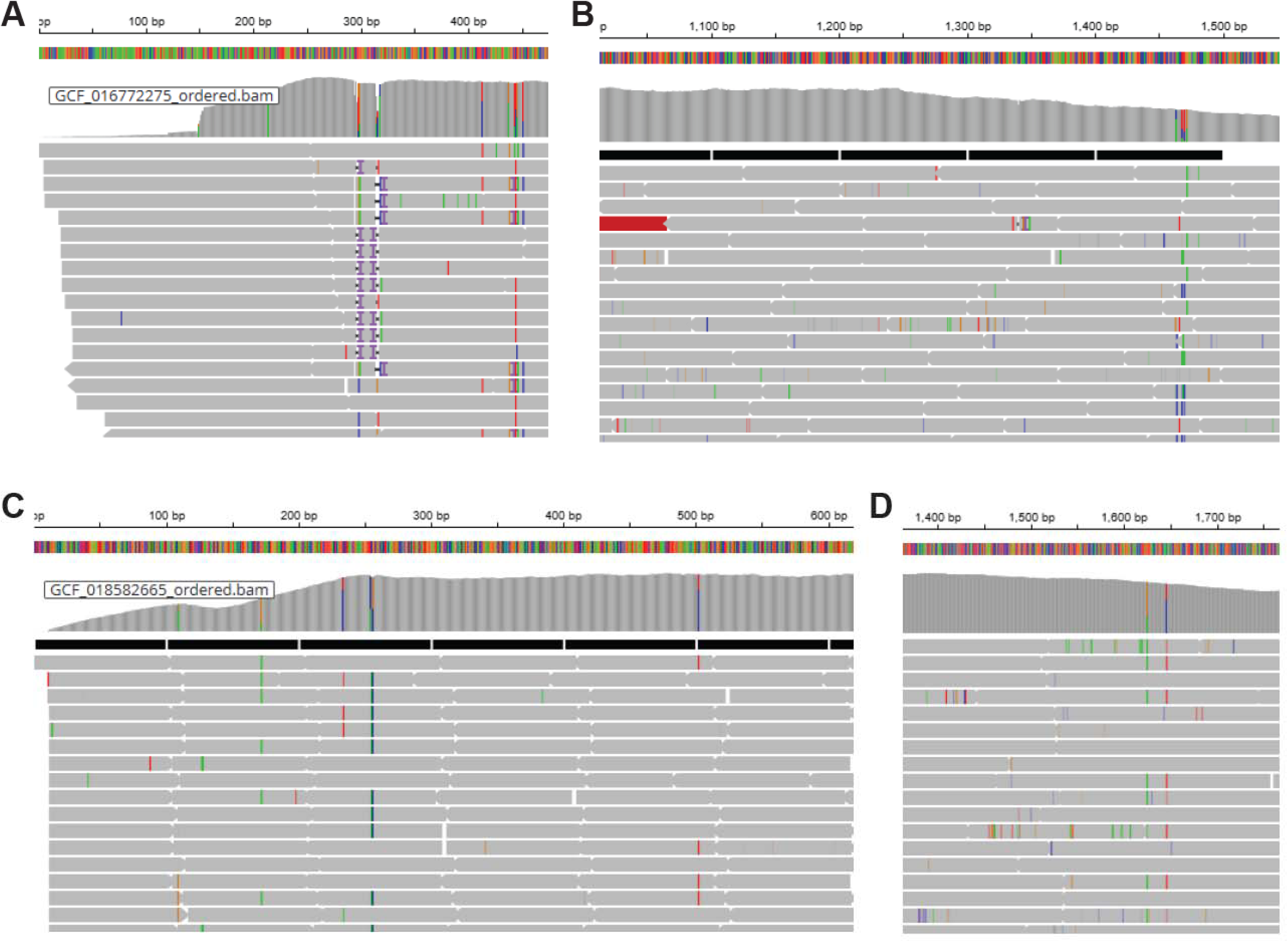
Biological variations in *16S rRNA* sequences of closed Bacillales genomes. Despite indicating one *16S rRN A* haplotype in the genome assembly, SyFi rightly finds multiple haplotypes clearly indicated by the presence of biological variations, either at the beginning of the sequence (A and C – GCF_016777275 and GCF_018582665) or at the end (B and D GCF_030123445 and GCF_004124315). All panels are snapshots from IGV.

**Figure S5.**
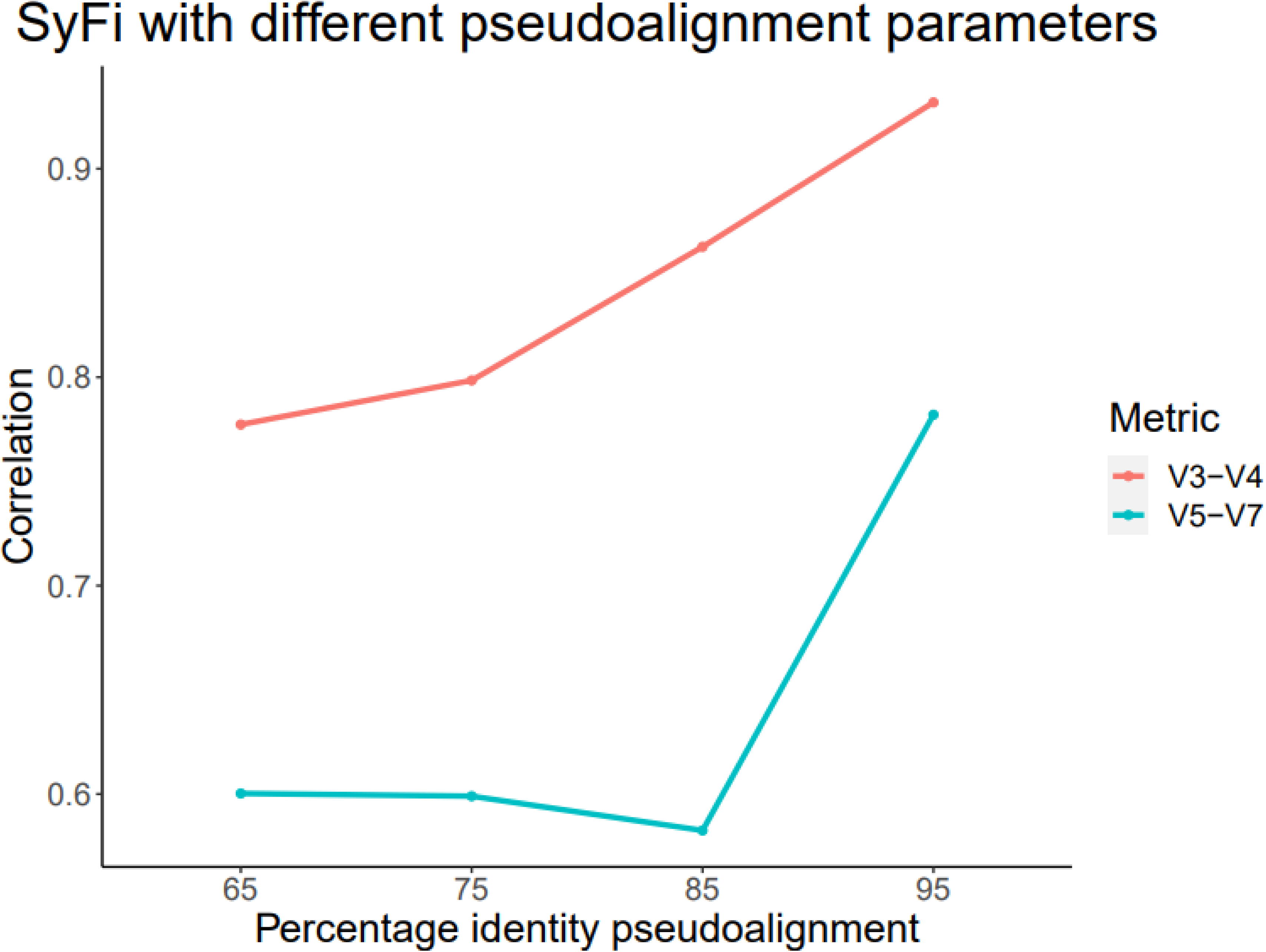
SyFi’s accuracy with different sequence percentage identity in pseudoalignment of metagenomic reads to *16S rRNA* V3-V4 or V5-V7 fingerprints. SyFi’s accuracy is assessed by the correlation to the shotgun metagenome dataset (y-axis) for each pseudoalignment percentage identity that is used in Salmon to match metagenomic reads with *16S rRNA* V3-V4 or V5- V7 fingerprints (x-axis).

**Figure S6.**
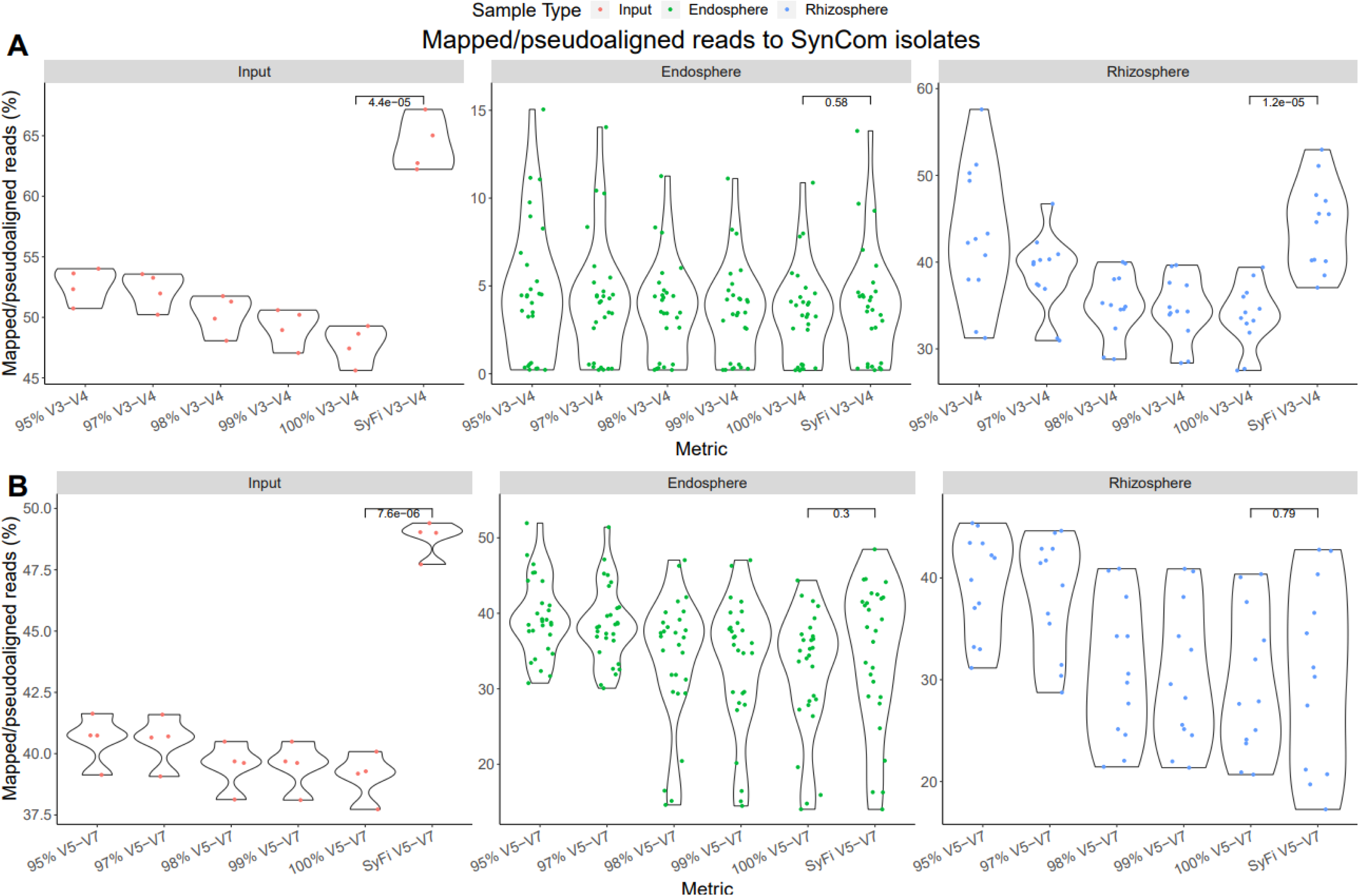
Percentages of mapped reads in the V3-V4 (A) and V5-V7 (B) datasets for microbial culture (Input), endosphere and rhizosphere samples. The relative amount of reads that are mapped to amplicon sequences using VSEARCH at different clustering percentage identities is compared to the relative amount of reads that Salmon is able to pseudoalign to the SyFi fingerprints. Statistical differences between SyFi and VSEARCH with 100% sequence identity are indicated by a pairwise t-test.

**Figure S7.**
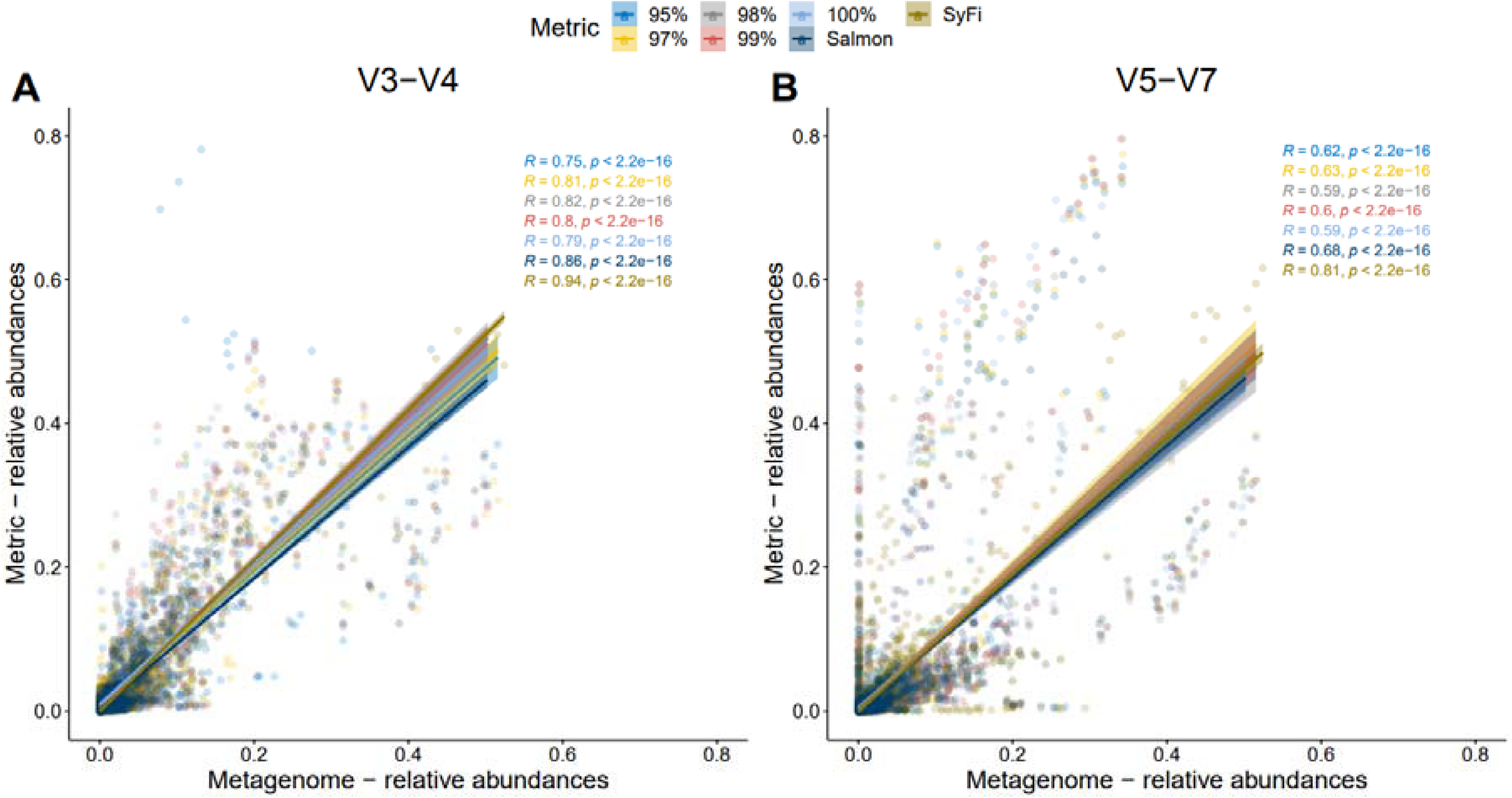
VSEARCH, Salmon and SyFi mapping accuracy to the metagenome dataset. Each coordinate is the relativ abundance of an isolate in a sample in the metagenome dataset (x-axis) and in the VSEARCH, Salmon or SyFi dataset (y-axis) for the V3-V4 dataset (A) and V5-V7 dataset (B). The Pearson correlation is shown in the top right corner with the p-value where the colors correspond to the dataset for each VSEARCH clustering sequence identity, for Salmon, pseudoaligning metagenomic reads to the amplicon sequences, or for SyFi, pseudoaligning metagenomic reads to the 16S rRNA V3-V4 or V5-V7 fingerprints.

## Supplementary tables

**Table S1.**
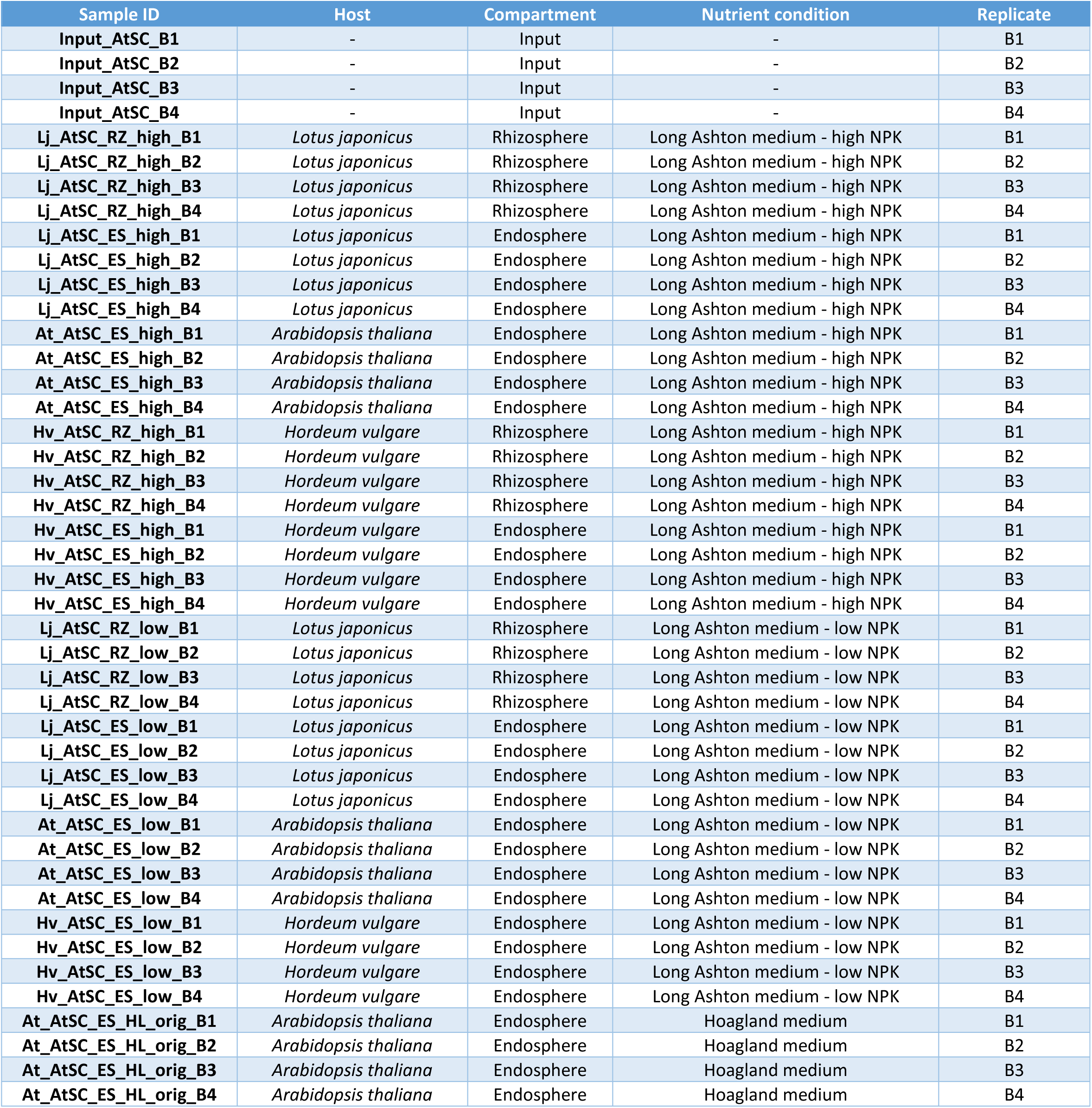
Sample overview for the SyFi validation, derived from Selten et al. (2024a).

**Table S2.**
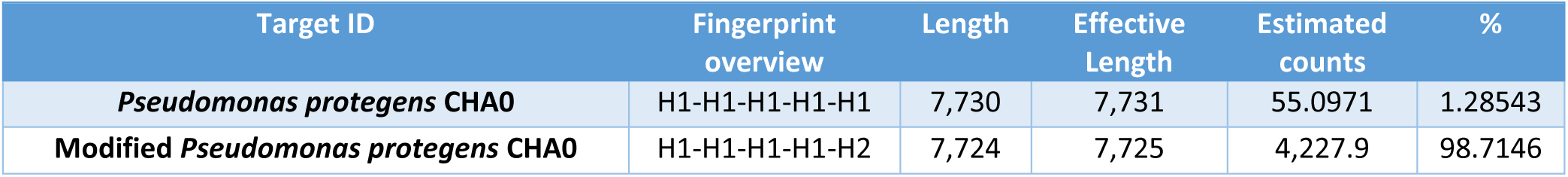
Explanation of pseudoalignment for distinguishing highly similar sequences. When pseudoaligning genomic reads from a computationally modified Pseudomonas protegens CHA0 to the wildtype strain and the modified strain, we find 98.7% of the reads to be correctly pseudoaligned.

**Table S3.** VSEARCH Clustering of V3-V4 and V5-V7 amplicon sequences and SyFi fingerprints on the plant root bacterial isolate dataset. The rows indicate whether VSEARCH was run on the original sequences that were retrieved directly from the genome clustered at different percentage identities (95, 97, 98, 99, 100% sequence identity) or on the SyFi fingerprints clustered at 100% sequence identity.

| VSEARCH clustering percentage (%) | Metric | No. of clusters | No. of singletons | No. of isolates in largest cluster | Average no. of isolates in cluster | No. of isolates |
| --- | --- | --- | --- | --- | --- | --- |
| **95** | V3-V4 | 39 | 10 | 62 | 9.8 | **382** |
| **97** | V3-V4 | 56 | 18 | 51 | 6.8 | **382** |
| **98** | V3-V4 | 66 | 22 | 41 | 5.8 | **382** |
| **99** | V3-V4 | 88 | 39 | 33 | 4.3 | **382** |
| **100** | V3-V4 | 136 | 77 | **23** | 2.8 | **382** |
| **SyFi** | V3-V4 | **161** | **102** | 24 | **2.4** | **382** |
| **95** | V5-V7 | 48 | 18 | 62 | 8 | **382** |
| **97** | V5-V7 | 62 | 24 | 62 | 6.2 | **382** |
| **98** | V5-V7 | 76 | 35 | 62 | 5 | **382** |
| **99** | V5-V7 | 95 | 47 | 48 | 4 | **382** |
| **100** | V5-V7 | 140 | 85 | 29 | 2.7 | **382** |
| **SyFi** | V5-V7 | **157** | **100** | **27** | **2.4** | **382** |

**Table S4.**
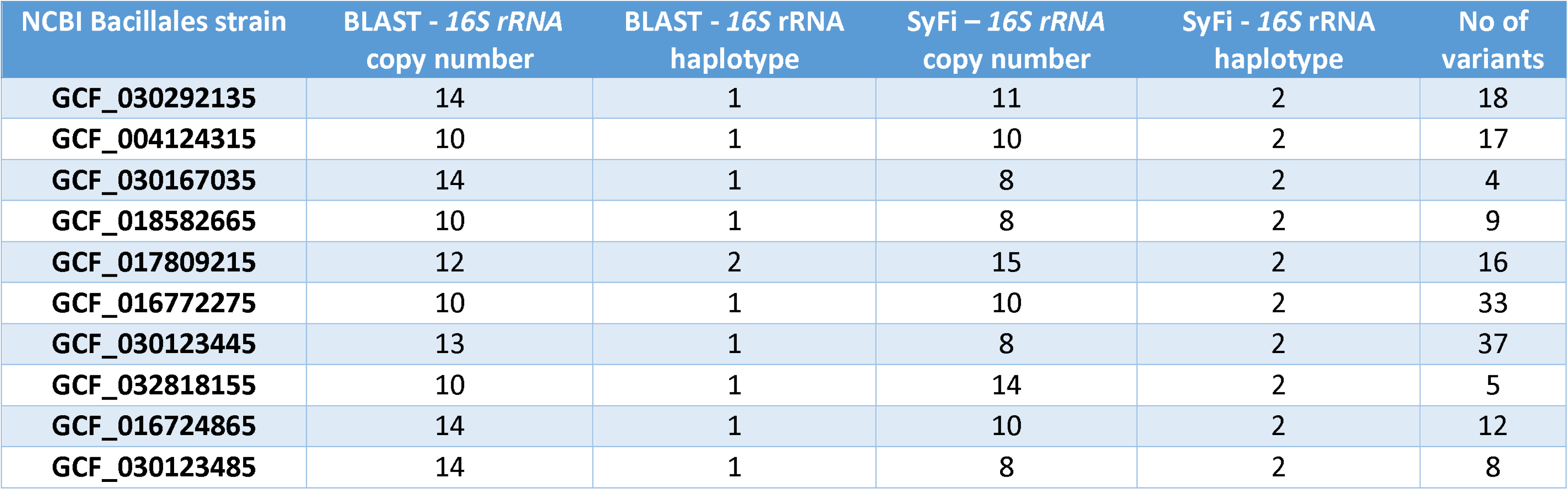
16S rRNA copy number and haplotype number in ten Bacillales NCBI complete genomes. These numbers are either found by direct target alignment of the 16S rRNA sequence of Paenisporosarcina (Supplemental sequence S1) to the genomes or by running SyFi on these genomes. Evidently SyFi finds more 16S rRNA haplotypes than BLAST, also illustrated by the number of variants found among the 16S rRNA copies.

| NCBI Bacillales strain | BLAST - 16S rRNA<br>copy number | BLAST - 16S rRNA<br>haplotype | SyFi – 16S rRNA<br>copy number | SyFi - 16S rRNA<br>haplotype | No of<br>variants |
| --- | --- | --- | --- | --- | --- |
| GCF_030292135 | 14 | 1 | 11 | 2 | 18 |
| GCF_004124315 | 10 | 1 | 10 | 2 | 17 |
| GCF_030167035 | 14 | 1 | 8 | 2 | 4 |
| GCF_018582665 | 10 | 1 | 8 | 2 | 9 |
| GCF_017809215 | 12 | 2 | 15 | 2 | 16 |
| GCF_016772275 | 10 | 1 | 10 | 2 | 33 |
| GCF_030123445 | 13 | 1 | 8 | 2 | 37 |
| GCF_032818155 | 10 | 1 | 14 | 2 | 5 |
| GCF_016724865 | 14 | 1 | 10 | 2 | 12 |
| GCF_030123485 | 14 | 1 | 8 | 2 | 8 |

**Table S5.** SyFi’s accuracy in assessment of presence/absence of isolates. The values indicate the percentage of isolates that were correctly identified as present or absent across all samples. The last columns are the sum of the correctly and incorrectly assessed percentages.

assessed percentages.
| Metric | Correctly<br>present | Incorrectly<br>present | Correctly<br>absent | Incorrectly<br>absent | Correct | Incorrect |
| --- | --- | --- | --- | --- | --- | --- |
| 95% V3-V4 | 22.8% | 2.0% | 63.9% | 11.3% | 86.7% | 13.3% |
| 97% V3-V4 | 18.8% | 2.0% | 69.8% | 9.4% | 88.6% | 11.4% |
| 98% V3-V4 | 18.8% | 1.1% | 72.0% | 8.1% | 90.7% | 9.3% |
| 99% V3-V4 | 14.4% | 1.0% | 76.7% | 8.0% | 91.1% | 8.9% |
| 100% V3-V4 | 9.7% | 1.1% | 81.1% | 8.1% | 90.8% | 9.2% |
| Salmon V3-V4 | 13.5% | 2.4% | 78.5% | 5.5% | 92.0% | 8.0% |
| SyFi V3-V4 | 10.3% | 2.1% | 82.5% | 5.1% | 92.8% | 7.2% |
| 95% V5-V7 | 20.9% | 2.5% | 68.0% | 8.6% | 88.9% | 11.1% |
| 97% V5-V7 | 16.7% | 1.9% | 74.2% | 7.3% | 90.9% | 9.1% |
| 98% V5-V7 | 17.0% | 0.7% | 77.2% | 5.1% | 94.2% | 5.8% |
| 99% V5-V7 | 14.0% | 1.0% | 79.8% | 5.2% | 93.8% | 6.2% |
| 100% V5-V7 | 11.3% | 1.4% | 83.1% | 4.2% | 94.4% | 5.6% |
| Salmon V5-V7 | 10.9% | 2.1% | 80.2% | 6.9% | 91.0% | 9.0% |
| SyFi V5-V7 | 8.9% | 2.2% | 83.1% | 5.7% | 92.1% | 7.9% |

## Supplementary sequence S1

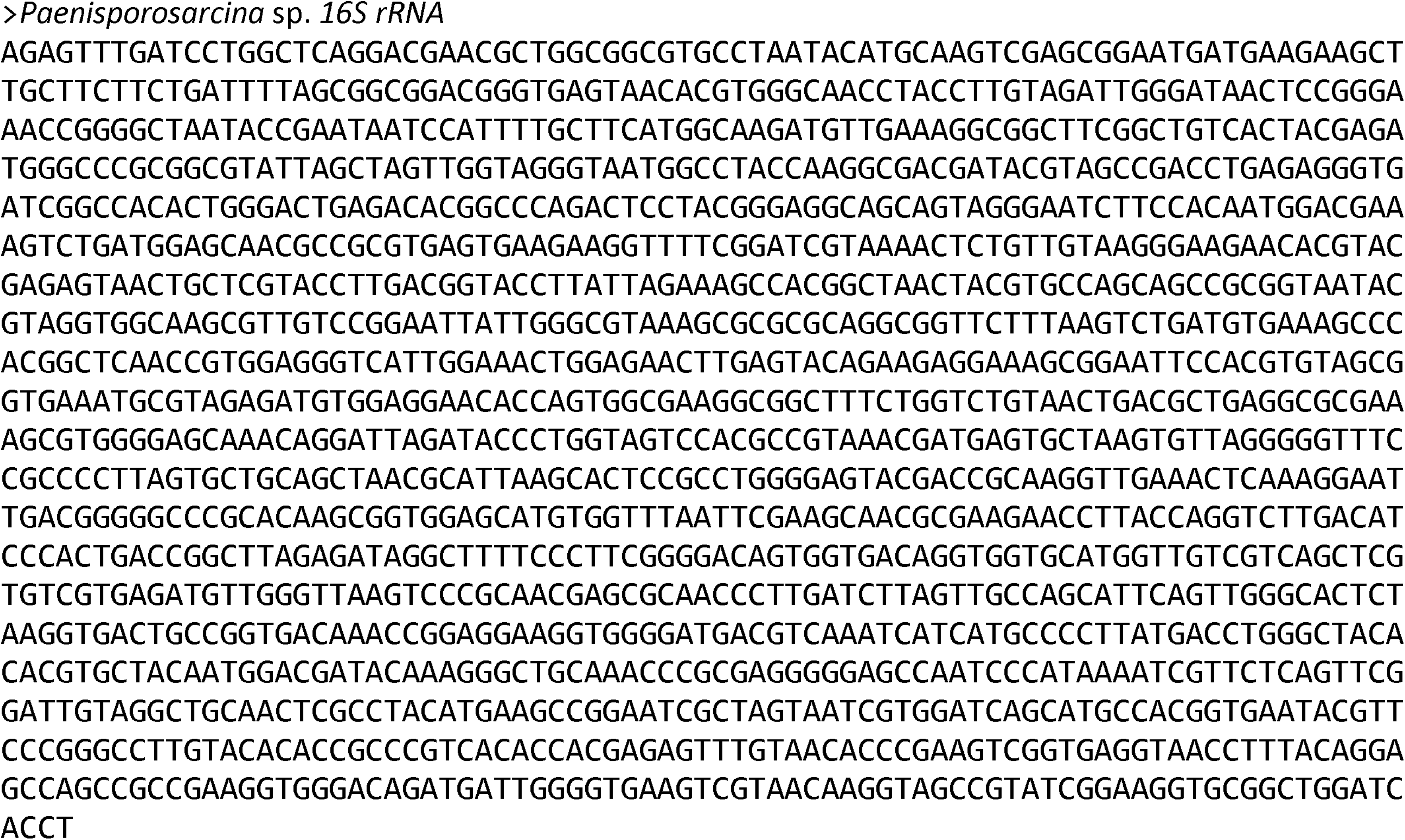

## Notes

### Competing Interest Statement

The authors have declared no competing interest.

