## Appendix for "SyFi: generating and using sequence fingerprints to distinguish SynCom isolates"

### Appendix 1 – *SyFi main* workflow

In *SyFi main*, a fingerprint of the target sequence for every bacterial genome is built (Figure 1). For this SyFi requires the genome sequence (fasta format), genomic single or paired end reads (fastq format), and a target sequence (fasta format) as input. *SyFi main*'s construction of the fingerprint can be divided into three steps.

In the first step, SyFi aligns the target sequence with the bacterial genome in a nucleotide-nucleotide BLAST alignment (v2.13.0) (default parameters) (Altschul et al., 1990) and extracts the nucleotide sequence with the highest identity score. Subsequently, the genomic reads are mapped to this sequence using BWA with default parameters (v2.2.1) (Li & Durbin, 2009) and filtered to obtain all the corresponding read pairs using Samtools (v1.16.1) (Li et al., 2009). The target sequence is then reassembled using SPAdes with default parameters (v3.15.5) (Bankevich et al., 2012) as SPAdes does not allow ambiguous bases in the target assembly. Biological variations, like SNPs, are masked by ambiguous nucleotides and should therefore be avoided. These SPAdes-assembled sequences are subsequently subjected to a length threshold to filter out sequences that are too small, after which the target sequence reads are filtered and trimmed according to the target sequence length by Samtools (v1.16.1) (Li et al., 2009). These steps are added to gain a clean target sequence from the genome with clean corresponding genomic reads.

In the second step, SyFi uses the target sequence and corresponding reads to call biological variations in the target sequence using the Picard algorithm (v2.27.5) in the GATK software (v3.8) (McKenna et al., 2010). After calling the variants in the target sequence, SyFi will enter three different modes depending on whether biological variations were found in the target sequence. When the variant calling workflow finds variants in the target sequence, the workflow will enter 'Mode 1' (Figure 1). If no variants are found, but SPAdes was already able to build multiple haplotypes, the target sequences will directly be forwarded to the Kallisto step in 'Mode 2'. When there is only one target sequence with no variants after the variant calling workflow, it indicates the bacterial isolate has only one haplotype. SyFi will then proceed in 'Mode 3' and calculate the copy number of this one haplotype before creating the fingerprint.

In Mode 1, Whatshap is used to identify biological variations that co-occur in the target sequence, which may indicate different variants, or haplotypes, of the target sequence (v1.7) (Martin et al., 2016). If there remain any identical haplotypes or haplotypes that are smaller than the length threshold, these are removed after the Whatshap step.

The third and last step of SyFi is to construct the fingerprint from the discovered haplotypes. In the case of Mode 1 and Mode 2, the transcript quantifying tool Kallisto (v0.48.0) (Bray et al., 2016) (default parameters) is employed to calculate the coverage of each target gene haplotype within the genome. This is calculated by pseudoalignment of genomic reads to the different haplotype sequences. In the meantime, the target gene copy number is estimated by comparing the target gene coverage to the genomic coverage. Together with the individual haplotype coverage values, these are used to estimate the occurrence of each haplotype in the bacterial genome. In the case of Mode 3, the target gene copy number indicates directly the occurrence of the only found haplotype.

In this copy number estimation, SyFi implements a threshold for the maximum number of copies. This can lead to the removal of haplotypes that occur in extremely low frequencies, which are likely biological contaminations or technical artifacts, either stemming from Whatshap (biological variations that might not occur together) or from sequencing errors. Table 1 shows an example on how the copy number of target sequences and haplotypes are estimated. Finally, the creation of a unique fingerprint is achieved by concatenating all the haplotypes into a fasta file according to their occurrences.

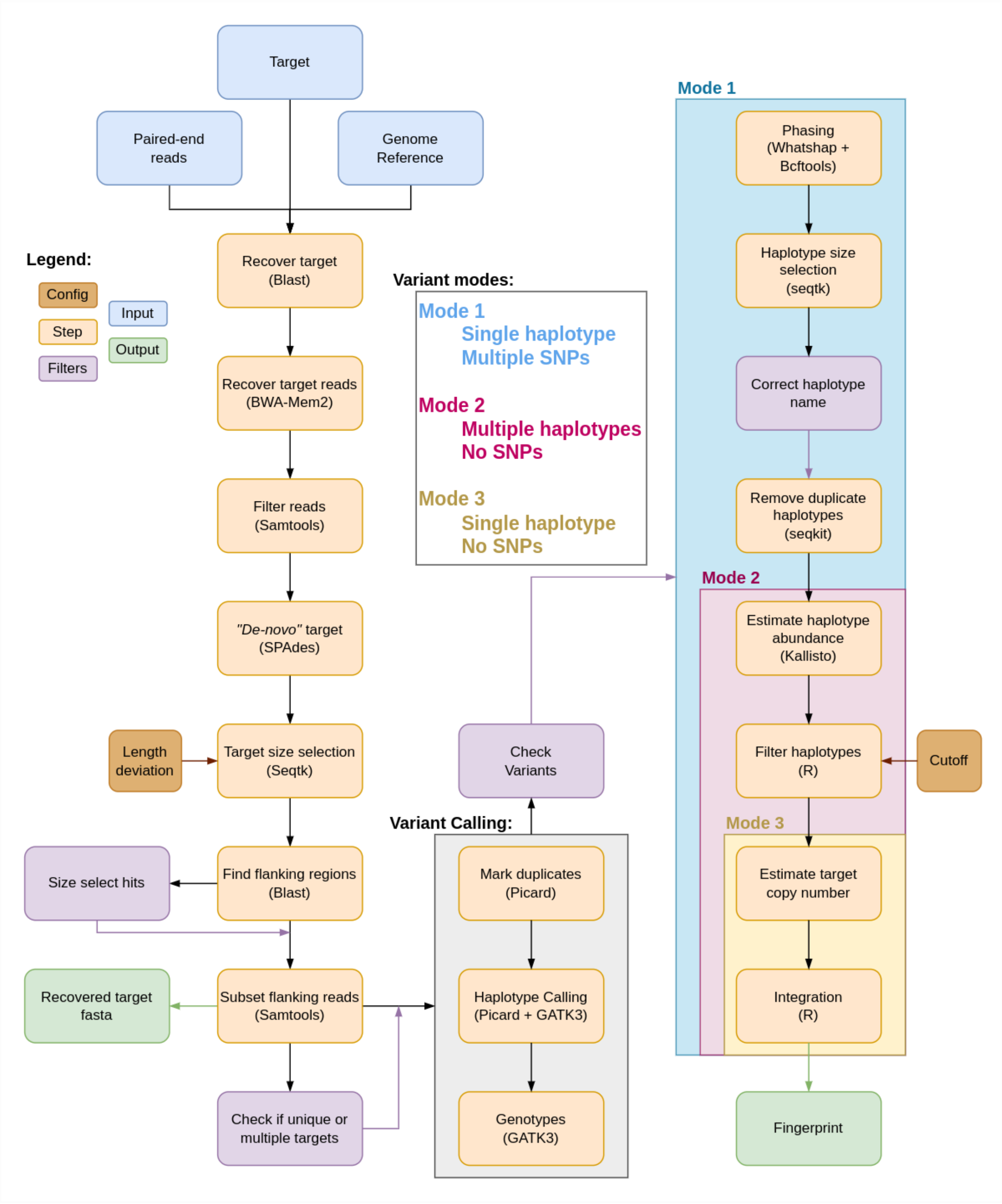


**Figure 1. Detailed overview of *SyFi main*. See text under Appendix 1 for clear explanation of all the steps.**

**Table 1. Building fingerprints.** The target sequence copy number is calculated by dividing the target coverage by the genome coverage (top). This copy number is subsequently used to estimate the haplotype occurrences in the bacterial genome (bottom)

| Strain | Genome length | Genome no of bases | Target length | Target no of bases | Genome coverage | Target Coverage | *16S rRNA* copy number | Rounded *16S rRNA* copy number |
|---|---|---|---|---|---|---|---|---|
| **KB_12** | 6249593 | 343125462 | 1511 | 328250 | 54.90365 | 217.2402 | 3.956754 | 4 |

| Haplotype | Target length | No of pseudoaligned reads | Ratio | rounded ratio | *16S rRNA* copy number | Occurrence |
|---|---|---|---|---|---|---|
| **H1** | 1511 | 1256.7 | 3.459172 | 3 | 4 | 3 |
| **H2** | 1511 | 363.295 | 1 | 1 | 4 | 1 |

#### Appendix 2 – SynCom data library preparation

To generate data for the SynCom reconstitution experiment, the *16S rRNA* V3-V4 and V5-V7 regions were subjected to high throughput two-step barcoding, purified using the AMPure XP Reagent for PCR purification (Beckman Coulter), purified for plant organelle-derived *16S rRNA* V5-V7 sequences, pooled and sequenced using a Novaseq 6000 PE250 sequencing run (2 x 250bp) (Table 1 and 2). The description of the shotgun metagenome-sequenced dataset can be found in (Selten et al., 2024).

**Table 1. PCR1 primers for *16S rRNA* V3-V4 and V5-V7 sequencing (top) and thermocycler settings of PCR 1 (bottom).**

| Primer name | Oligo sequence |
|---|---|
| ***16S rRNA* V5-V7 F** | TCGTCGGCAGCGTCAGATGTGTATAAGAGACAGAACMGGATTAGATACCCKG |
| ***16S rRNA* V5-V7 R** | GTCTCGTGGGCTCGGAGATGTGTATAAGAGACAGACGTCATCCCCACCTTCC |
| ***16S rRNA* V3-V4 F** | TCGTCGGCAGCGTCAGATGTGTATAAGAGACAGCCTACGGGNGGCWGCAG |
| ***16S rRNA* V3-V4 R** | GTCTCGTGGGCTCGGAGATGTGTATAAGAGACAGGACTACHVGGGTATCTAATCC |

| PCR1 settings | T (°C) | Time | |
|---|---|---|---|
| **First denaturation** | 95 | 2 min | |
| **Denaturation** | 95 | 30 s | x30 cycles |
| **Annealing** | 55 | 30 s | |
| **Elongation** | 72 | 45 s | |
| **Final elongation** | 72 | 10 min | |
| **End** | 12 | ∞ | |

**Table 2. PCR2 primers for V3-V4 and V5-V7 *16S rRNA* sequencing and PCR2 thermocycler settings. This step consisted of one universal forward primer and barcoded reverse primers for amplification**

| | Primer name | Oligo sequence |
|---|---|---|
| V3-V4 | **UDP0200-i5** | AATGATACGGCGACCACCGAGATCTACACAATGTATTGCTCGTCGGCAGCGTC |
| V5-V7 | **UDP0201-i5** | AATGATACGGCGACCACCGAGATCTACACGATCTCTGGATCGTCGGCAGCGTC |
| | *UDP0289V2-i7* | CAAGCAGAAGACGGCATACGAGATGCTACTATCTGTCTCGTGGGCTCGG |
| | *UDP0289-i7* | CAAGCAGAAGACGGCATACGAGATGGAATTGTTCGTCTCGTGGGCTCGG |
| | *UDP0290V2-i7* | CAAGCAGAAGACGGCATACGAGATGTCTTCTAATGTCTCGTGGGCTCGG |
| | *UDP0290-i7* | CAAGCAGAAGACGGCATACGAGATCCGGACCACAGTCTCGTGGGCTCGG |
| | *UDP0291V2-i7* | CAAGCAGAAGACGGCATACGAGATATGTGCGAGCGTCTCGTGGGCTCGG |
| | *UDP0291-i7* | CAAGCAGAAGACGGCATACGAGATGACTTAGAAGGTCTCGTGGGCTCGG |
| | *UDP0292-i7* | CAAGCAGAAGACGGCATACGAGATTGGCAATATTGTCTCGTGGGCTCGG |
| | *UDP0293-i7* | CAAGCAGAAGACGGCATACGAGATGAATGCACGAGTCTCGTGGGCTCGG |
| | *UDP0294-i7* | CAAGCAGAAGACGGCATACGAGATCGTGTATCTTGTCTCGTGGGCTCGG |
| | *UDP0295-i7* | CAAGCAGAAGACGGCATACGAGATATTCATTGCAGTCTCGTGGGCTCGG |
| | *UDP0296-i7* | CAAGCAGAAGACGGCATACGAGATTCCTTCATAGGTCTCGTGGGCTCGG |
| | *UDP0297-i7* | CAAGCAGAAGACGGCATACGAGATTCTAGTCTTCGTCTCGTGGGCTCGG |
| | *UDP0298-i7* | CAAGCAGAAGACGGCATACGAGATCTCGACTCCTGTCTCGTGGGCTCGG |
| | *UDP0299-i7* | CAAGCAGAAGACGGCATACGAGATAGTGAGTGAAGTCTCGTGGGCTCGG |

| | | |
|---|---|---|
| | *UDP0300-i7* | CAAGCAGAAGACGGCATACGAGATGAAGCGGACCGTCTCGTGGGCTCGG |
| | *UDP0301V2-i7* | CAAGCAGAAGACGGCATACGAGATCAAGCCACTAGTCTCGTGGGCTCGG |
| | *UDP0301-i7* | CAAGCAGAAGACGGCATACGAGATGCTCTCGTTGGTCTCGTGGGCTCGG |
| | *UDP0302-i7* | CAAGCAGAAGACGGCATACGAGATGGACCTCAATGTCTCGTGGGCTCGG |
| | *UDP0303-i7* | CAAGCAGAAGACGGCATACGAGATGAGTCTCTCCGTCTCGTGGGCTCGG |
| | *UDP0304-i7* | CAAGCAGAAGACGGCATACGAGATAACGGAGCGGGTCTCGTGGGCTCGG |
| | *UDP0305-i7* | CAAGCAGAAGACGGCATACGAGATTGTGATGTATGTCTCGTGGGCTCGG |
| | *UDP0306-i7* | CAAGCAGAAGACGGCATACGAGATAACATACCTAGTCTCGTGGGCTCGG |
| | *UDP0307-i7* | CAAGCAGAAGACGGCATACGAGATGTGCTAGGTGGTCTCGTGGGCTCGG |
| | *UDP0308-i7* | CAAGCAGAAGACGGCATACGAGATCATACTTGAAGTCTCGTGGGCTCGG |
| | *UDP0309-i7* | CAAGCAGAAGACGGCATACGAGATCTTGTCTTAAGTCTCGTGGGCTCGG |
| | *UDP0310-i7* | CAAGCAGAAGACGGCATACGAGATAAGAGAGGTGGTCTCGTGGGCTCGG |
| | *UDP0311-i7* | CAAGCAGAAGACGGCATACGAGATTGCACGAGAAGTCTCGTGGGCTCGG |
| | *UDP0312-i7* | CAAGCAGAAGACGGCATACGAGATACTTCCTAGCGTCTCGTGGGCTCGG |
| | *UDP0313-i7* | CAAGCAGAAGACGGCATACGAGATGTGCTATTAAGTCTCGTGGGCTCGG |
| | *UDP0314-i7* | CAAGCAGAAGACGGCATACGAGATAGCGTGAATGGTCTCGTGGGCTCGG |
| | *UDP0315-i7* | CAAGCAGAAGACGGCATACGAGATCCTTAGTGCCGTCTCGTGGGCTCGG |
| | *UDP0316-i7* | CAAGCAGAAGACGGCATACGAGATTGTACCGAATGTCTCGTGGGCTCGG |
| | *UDP0317-i7* | CAAGCAGAAGACGGCATACGAGATGGAGATTAGTGTCTCGTGGGCTCGG |
| | *UDP0318-i7* | CAAGCAGAAGACGGCATACGAGATTACTAACACAGTCTCGTGGGCTCGG |
| | *UDP0319-i7* | CAAGCAGAAGACGGCATACGAGATTAGGTCGTTGGTCTCGTGGGCTCGG |
| | *UDP0320-i7* | CAAGCAGAAGACGGCATACGAGATATGCCGACCGGTCTCGTGGGCTCGG |
| | *UDP0321-i7* | CAAGCAGAAGACGGCATACGAGATCTAGCGTCGAGTCTCGTGGGCTCGG |
| | *UDP0322-i7* | CAAGCAGAAGACGGCATACGAGATTGCCTACGAGGTCTCGTGGGCTCGG |
| | *UDP0323-i7* | CAAGCAGAAGACGGCATACGAGATACTAGAACTTGTCTCGTGGGCTCGG |
| | *UDP0324-i7* | CAAGCAGAAGACGGCATACGAGATCACCTCTTGGGTCTCGTGGGCTCGG |
| | *UDP0325-i7* | CAAGCAGAAGACGGCATACGAGATAAGCAGATATGTCTCGTGGGCTCGG |
| | *UDP0326-i7* | CAAGCAGAAGACGGCATACGAGATGCCAGATCCAGTCTCGTGGGCTCGG |
| | *UDP0327-i7* | CAAGCAGAAGACGGCATACGAGATTTGGATTCAAGTCTCGTGGGCTCGG |

| PCR 2 settings | T°C | Time | |
|---|---|---|---|
| **First denaturation** | 95 | 2 min | |
| **Denaturation** | 95 | 30 s | x10 cycles |
| **Annealing** | 55 | 30 s | |
| **Elongation** | 72 | 45 s | |
| **Final elongation** | 72 | 10 min | |
| **End** | 12 | ∞ | |

#### Appendix 3 – Contamination, heterogeneity, and GC content affects determination of *16S rRNA* copy number

Several of the 447 plant root bacterial isolates are indicated by exceptionally high *16S rRNA* copy numbers, with more than twenty *16S rRNA* sequences found in these genomes (Manuscript: Figure 3A and C). Interestingly, these high *16S rRNA* copy numbers are also observed in a few complete NCBI genomes (Manuscript: Figure S3A and S4). To investigate whether these high *16S rRNA* copy numbers are real and not technical artifacts, we correlated the *16S rRNA* copy number with the level of contamination and heterogeneity in the genome assemblies that we derived from CheckM (version 1.1.3) (Parks et al., 2015). Contamination and heterogeneity occur when single-copy marker genes are found more than once, either due to contamination from another bacterium (contamination) or from a duplication event (heterogeneity). In addition, a contamination from a phylogenetically closely related strain is often recognized as heterogeneity as well. Evidently, we discovered a weak correlation between the *16S rRNA* copy number and contamination level and no correlation with the heterogeneity level (Figure 1A). Conclusively, *16S rRNA* copies from contaminating sequences does not explain the exceptionally high *16S rRNA* copy numbers, though may play role for a couple of genomes.

Another explanation for the exceptionally high number of *16S rRNA* copies could be related to the GC content. Whole-genome sequencing of bacterial strains does not result in uniform read distribution or coverage across the entire genome, as GC-rich regions can experience higher coverage as compared to GC-poor regions (Chen et al., 2013; Tyler et al., 2016; Gunasekera et al., 2021). To investigate this, we correlated the GC content of the *16S rRNA* fingerprints to the *16S rRNA* copy number (Figure 1B). We also compared the GC content of the *16S rRNA* sequence from the complete NCBI genomes with their *16S rRNA* copy number to assess to what extent GC content correlates with overestimation of *16S rRNA* copy number (Figure 1C). Indeed, we found a significant positive correlation between the GC content of the *16S rRNA* fingerprint or sequence and the *16S rRNA* copy number in both datasets (Figure 1B and C). By comparing the correlation slopes between the two datasets we can estimate the extent to which SyFi may overestimate the *16S rRNA* copy number compared to copy numbers of closed genomes, assuming that fully assembled genomes did not lose any *16S rRNA* copies. The difference in correlation slopes indicates how SyFi might be extensively overestimating the *16S rRNA* copy number at high GC contents in the bacterial genome (Figure 1D black line). This observation, however, may also potentially differ between different bacterial taxonomic groups, corroborating the need for a phylogenetic marker gene database in future implementations of SyFi.

Conclusively, contamination may contribute to high, biologically infeasible *16S rRNA* copy numbers. In spite of this finding, a high GC content seems mainly responsible for *16S rRNA* copy number overestimation. In future versions, a GC content normalization of the *16S rRNA* copy number may be implemented in SyFi to account for this overestimation.

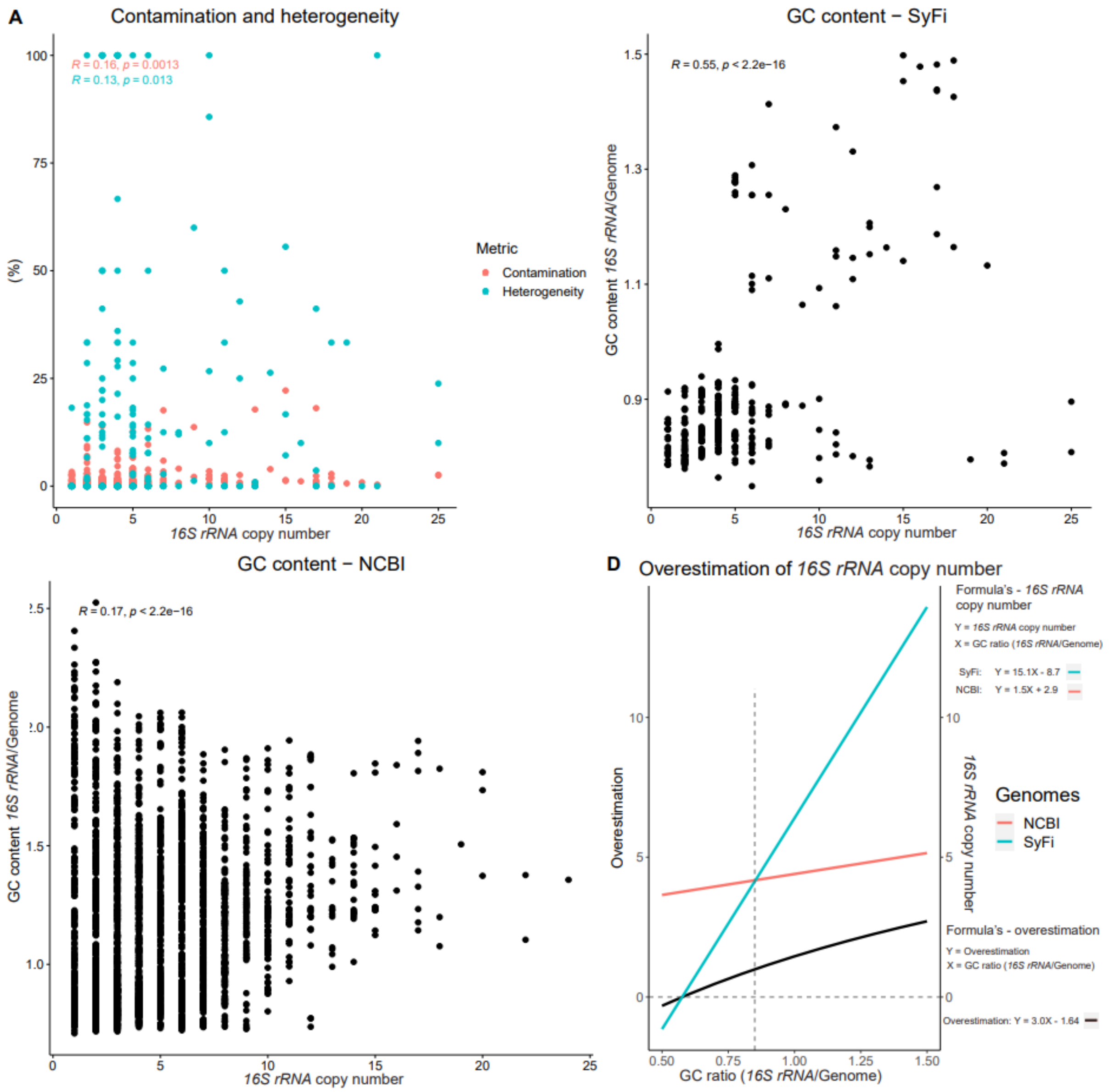


**Figure 1. The effect of contamination, heterogeneity, and GC content on the *16S rRNA* copy.** Contamination and heterogeneity (A) (y-axis) in the genome assemblies correlate weakly but significantly with the *16S rRNA* copy number (x-axis). A significant positive correlation exists between proportion of GC content in the *16S rRNA* sequence as compared to the genome (B) (y-axis) and the *16S rRNA* copy number (x-axis). The *16S rRNA* copy number in complete NCBI genomes is also affected by the difference in GC content between the *16S rRNA* sequence and the genome (C). The correlation between *16S rRNA* copy number and *16S rRNA*-genome GC content among the genomes used in SyFi as well as the closed NCBI genomes is compared to each other to illustrate the overestimation of *16S rRNA* copy number (D).
